# CRISPR-Csy4-mediated editing of rotavirus double-stranded RNA genome

**DOI:** 10.1101/2020.03.09.983262

**Authors:** Guido Papa, Luca Venditti, Luca Braga, Edoardo Schneider, Mauro Giacca, Gianluca Petris, Oscar R. Burrone

## Abstract

CRISPR-nucleases have been widely applied for editing cellular and viral genomes, but nuclease-mediated genome editing of double-stranded RNA (dsRNA) viruses has not yet been reported. Here, by engineering CRISPR-Csy4 nuclease to localise to rotavirus viral factories, we achieved the first nuclease-mediated genome editing of rotavirus, an important human and livestock pathogen with a multi-segmented dsRNA genome. Rotavirus replication intermediates cleaved by Csy4 were repaired through the formation of defined deletions in the targeted genome segments in a single replication cycle. Using CRISPR-Csy4-mediated editing of rotavirus genome, we labelled for the first time the products of rotavirus secondary transcription made by newly assembled viral particles during rotavirus replication, demonstrating that this step largely contributes to the overall production of viral proteins. We anticipate that the nuclease-mediated cleavage of dsRNA virus genomes will promote a new level of understanding of viral replication and host-pathogen interactions, offering the opportunity to develop new therapeutics.

## INTRODUCTION

Prokaryotes have evolved an anti-viral defence mechanism, which relies on clustered, regularly interspaced, short palindromic repeat (CRISPR) loci. These regions contain short virus-derived sequences separated by conserved repeated sequences, which are transcribed as pre-CRISPR RNAs (pre-crRNAs). These transcripts are then processed to generate crRNAs, which are used to guide CRISPR adaptive immunity to cleave foreign nucleic acids (Makarova et al., 2018; Marraffini and Sontheimer, 2010). The six known types of CRISPR-Cas systems have different mechanisms of crRNA maturation (Makarova et al., 2018). In the type II CRISPR-Cas9 system, well known for its application in genome editing, RNAse III mediates maturation of gRNAs from the pre-crRNAs and trans-activating crRNA (tracrRNA). In CRISPR type V-A and type VI guide RNAs (gRNAs) are formed only by the crRNA, which is directly processed by the targeted nuclease Cas12a and Cas13, respectively (Makarova et al., 2018).

The CRISPR-Cas type I and type III (and likely type IV) systems, instead, rely on the use of an endoribonuclease from the Cas6 superfamily to cleave pre-crRNAs within each invariant repeat sequence to generate mature crRNAs (Murugan et al., 2017; Özcan et al., 2019). Among them, the Cas6 protein of *Pseudomonas aeruginosa* type I-F CRISPR systems, known as Csy4/Cas6f (Makarova et al., 2011, 2018), has been well-characterised as a small (21 kDa) and highly specific RNA endoribonuclease. Csy4 endoribonuclease processes, as a single-turnover enzyme, pre-crRNAs containing a 28-nucleotides near-identical repeats (Cy28) to generate the mature crRNAs. In the Cy28 sequence Csy4 specifically binds to the 16 nucleotides RNA hairpin with very high affinity (K_d_=50 pM) and cleaves directly downstream of the five-base-pair stem element (Haurwitz et al., 2012, 2010; Sternberg et al., 2012; Young et al., 2013). Because of all these very useful enzymatic features, Csy4 nuclease has already been applied to cleave several engineered Cy28-containing RNAs for biotechnological applications including a model HIV-1 virus (Guo et al., 2015; Nissim et al., 2014; Young et al., 2013).

In this study, we applied Csy4 on rotavirus (RV), an important animal and human pathogen of the *Reoviridae* family, with a multi-segmented dsRNA genome, which replicates in cytoplasmic phase-dense structures (viral factories) named viroplasms (Desselberger, 2014; Eichwald et al., 2004; Estes and Greenberg, 2013; Fabbretti et al., 1999). Protein access to viroplasms is mainly restricted to some viral proteins and is regulated in a still unknown manner (Eichwald et al., 2012). As a consequence, also exogenous non-viral proteins are usually confined outside these viral factories (Silvestri et al., 2004). In the past, we succeeded to localise EGFP and mCherry into viroplasms by fusing them to either the RV non-structural proteins NSP5 or NSP2 (Eichwald et al., 2004; Papa et al., 2019). Here, we further expand the use of these two proteins as a general shuttle to localise CRISPR-Csy4 nuclease to RV viroplasms and obtain the first nuclease-mediated site-specific genome editing of a dsRNA virus. Using different recombinant RVs (rRVs) carrying the Csy4 target sequence in diverse positions and viral segments, we report that nuclease cleavage of a viral RNA within viroplasms results in small sequence-dependent deletions of the targeted genomic segment in a single replication cycle. Thanks to RV genome editing, we now shed light on a still debated aspect of RV replication cycle. For the first time, we label and measure the products of the secondary transcription made by the newly assembled RV intermediate particles, demonstrating the main role of secondary transcription to the overall production of viral proteins in virus-infected cells.

## RESULTS

### Targeting CRISPR-Csy4 to RV viroplasms

To investigate the effect of Csy4 cleavage of RV plus strand RNAs, we first generated a panel of different Csy4 stable cell lines in MA104 cells, which are the most used cells to study rotavirus (Wu et al., 2017). We found that the SV5-tagged Csy4 was expressed at low levels (MA-Csy4) compared to the catalytically inactive H29A Csy4 variant (MA-Csy4-H29A) (Figure 1A), probably because of the reported Csy4 nuclease toxicity when expressed at high levels in eukaryotic cells (Nissim et al., 2014). As Csy4-H29A mutation preserves most Csy4 functions, including the strong binding affinity for the hairpin target sequence (Haurwitz et al., 2010; Sternberg et al., 2012), we took advantage of its higher expression to evaluate its localisation in wild-type recombinant RV (rRV-wt) infected and non-infected cells by immunofluorescence. Csy4-H29A showed broadly diffused pattern with no viroplasm localisation in both, non-infected and rRV-infected cells (Figure 1B). Considering that NSP5 and NSP2 are essential and highly abundant rotavirus non-structural proteins localising to viroplasms (Eichwald et al., 2004; Martin et al., 2011; Papa et al., 2019), we fused each of them to the SV5-tagged Csy4 nuclease and generated two MA104 stable cell lines, namely MA-NSP5-Csy4 and MA-NSP2-Csy4 (Figure 1C). As control, NSP5 was also fused to the SV5-tagged Csy4-H29A (MA-NSP5-Csy4-H29A) (Figure 1C). Both NSP5-Csy4 and NSP5-Csy4-H29A fusion proteins showed higher expression than the NSP2-Csy4 chimera (Figure 1C). All three fusion proteins efficiently localised to viroplasms upon RV infection, while they showed a diffused pattern in non-infected cells (Figure 1D-F). In MA-NSP5-Csy4 and MA-NSP5-Csy4-H29A cell lines the number and size of viroplasms, as well as production of viral proteins (VP2 and NSP5), did not differ from those in the parental MA104 cells (Figure 1; Figure S1A-C), indicating that the presence of these fusion proteins did not affect viroplasms formation.

**Figure 1.**
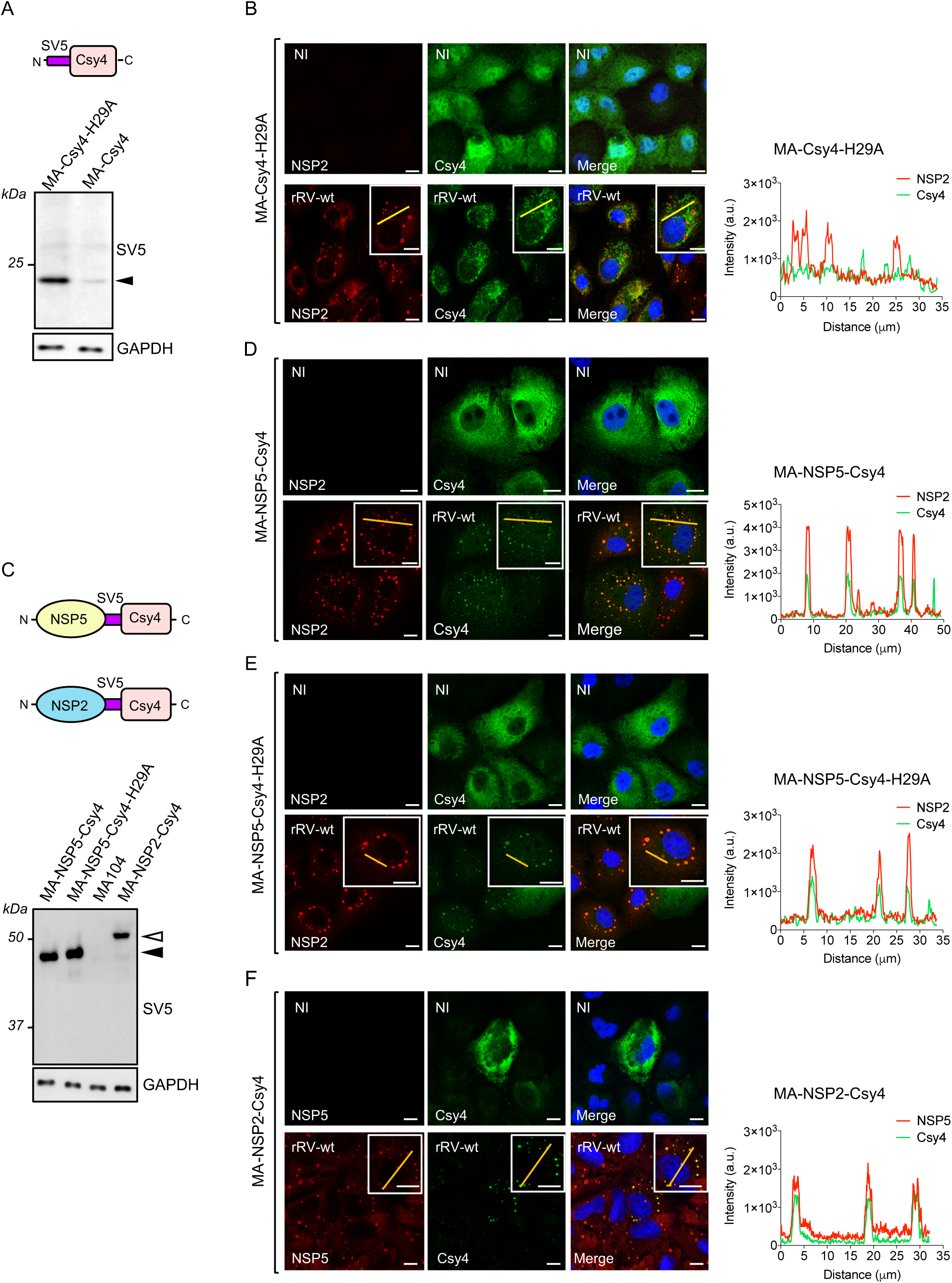
NSP5-Csy4 and NSP2-Csy4 localise to viroplasms in RV infected MA104 cells. A) Upper panel, scheme of the SV5-tagged Csy4 protein; lower panel, western blot of total cell extracts from SV5-tagged Csy4-H29A (MA-Csy4-H29A) and Csy4 (MA-Csy4) MA104 cell lines developed with anti-SV5. GAPDH was used as loading control. Black arrowhead indicates the position of SV5-Csy4 and SV5-Csy4-H29A. B) Immunofluorescence of rRV-wt infected (MOI of 10) and non-infected MA-Csy4-H29A cells. Csy4 was labelled with anti-SV5 (green), viroplasms with anti-NSP2 (red) and nuclei with DAPI (blue). Fluorescent intensity profile of rRV-wt infected MA-Csy4-H29A cells is shown on the right panel. Yellow line indicates area of measurement. C) Upper panel, scheme of the NSP5 and NSP2 SV5-Csy4 fusion proteins; lower panel, western blot of total cell extracts of the indicated stable transfectant cell lines and of MA104, developed with anti-SV5. Filled arrowhead indicates Csy4 fused to NSP5 and open arrowhead indicates Csy4 fused to NSP2. D-F) Immunofluorescence of rRV-wt infected and non-infected cell lines expressing NSP5-Csy4 (D), NSP5-Csy4-H29A (E) and NSP2-Csy4 cells (F). Fluorescent intensity profiles of the indicated cell line infected with rRV-wt is shown on the right panel of (D), (E), (F). Csy4 fusion proteins were labelled with anti-SV5 (green), viroplasms with anti-NSP2 (red in D and E) or anti-NSP5 (red in F) and nuclei with DAPI (blue). Scale bar 13 μm. In all panels data are representative of at least n=2 independent experiments.

### Activity of Csy4 and Csy4 fusion variants

RNA cleavage activity of the different Csy4 variants was evaluated using an EGFP reporter plasmid having the Cy28 target sequence located immediately after the ATG initiation codon (Figure 2A) (Borchardt et al., 2015). Csy4-mediated cleavage of the reporter transcript was 5 measured by the reduction of EGFP intensity (Borchardt et al., 2015). As shown in Figure 2A, Cy28-EGFP levels were strongly but similarly diminished in MA-NSP2-Csy4 and MA-Csy4 cells, compared to the high levels observed in cells expressing the inactive H29A mutant (MA-NSP5-Csy4-H29A, MA-Csy4-H29A) and the parental MA104, despite NSP2-Csy4 been expressed at higher levels than Csy4 (Figure1A, 1C). The most reduced Cy28-EGFP expression was, however, observed in MA-NSP5-Csy4 cells (Figure 2A), which expressed more Csy4 fusion compared to MA-NSP2-Csy4 cells (Figure 1C). Expression levels of control EGFP lacking the Cy28 target sequence showed no differences among all the samples (Figure 2B). These data suggest that although the NSP2/NSP5-Csy4 fusion proteins may not be as efficient as SV5-Csy4, they retain a similar activity thanks to their higher expression level, representing an ideal tool to evaluate the effect of Csy4 nuclease activity on RV genome replication.

**Figure 2.**
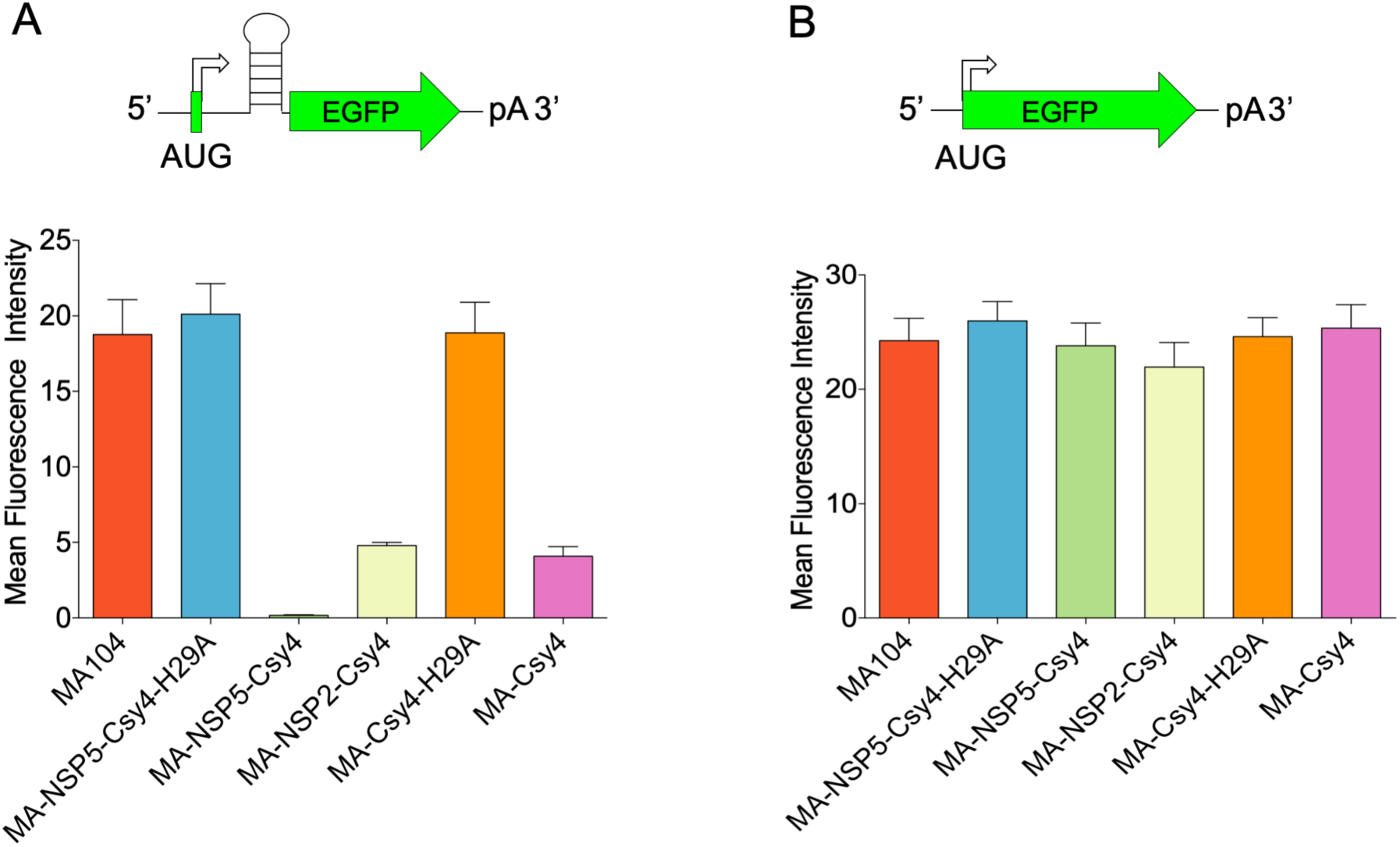
Activity of Csy4 and Csy4 fusion variants on the EGFP reporter model. A,B) Upper panels scheme or reporter EGFP transcripts; lower panels, EGFP fluorescence of the indicated cell lines transfected with pEGFP-Cy28 (A) or pEGFP (B) plasmids. Data are means ± SEM from n=3 independent experiments.

### Viroplasm-targeted Csy4 nuclease mediates editing of RV gs5

The RV genome segment 5 (gs5) includes a single open reading frame (ORF) encoding the 60 kDa non-structural viral protein NSP1, which functions as an antagonist of the interferon response (Barro and Patton, 2007; Davis and Patton, 2017). Despite its involvement in counteracting cellular immune system, several studies showed that expression of NSP1 is nevertheless dispensable for virus replication in cultured cells (Kanai et al., 2017, 2018; Komoto et al., 2018). We took advantage of the recently developed fully-tractable RV reverse genetics (RG) system (Komoto et al., 2018) to generate a recombinant Rotavirus (rRV) of the simian strain SA11 containing a modified version of gs5. We engineered gs5 (herein named gs5*) to contain the Cy28 in the positive RNA strand between the NSP1 STOP codon and its 3’ UnTranslated Region (3’UTR) (Figure 3A). NSP1 was also C-terminally tagged with SV5 to facilitate detection of the recombinant viral protein (Figure 3A). Analysis of the genomic dsRNA migration profile of the new recombinant Rotavirus, named rRV-gs5*, showed the expected increase in size of gs5 (gs5*, red arrow), also confirmed by sequencing, and packaging of all the other genome segments (Figure S2A-B). rRV-gs5*-infected MA104 cells produced the 60 kDa SV5-tagged NSP1 protein and no evident differences in the hyperphosphorylation of the RV essential protein NSP5 (Figure S2C). rRV-gs5* showed minimal changes in viral replication at early times post infection compared to the rRV with unmodified gs5 (rRV-wt), even though a slightly reduced replication fitness was present at late times post infection (Figure S2D). This may be due to a partial impairment of interferon antagonist activity for the C-terminally SV5-tagged NSP1 compared to the wild-type protein (Kanai et al., 2017).

**Figure 3.**
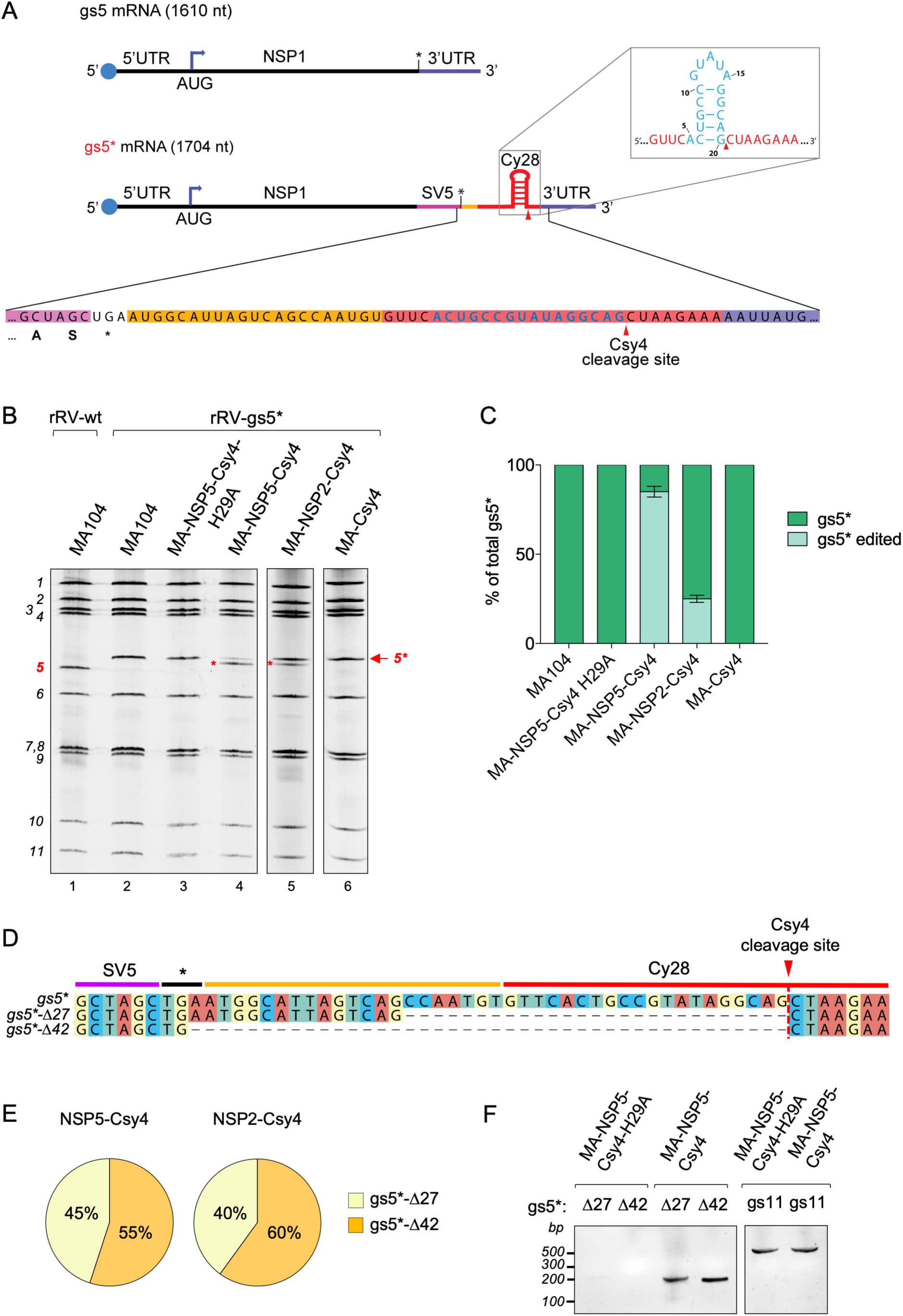
CRISPR-Csy4 editing of rRV genome segment 5*. A) Scheme of wild type gs5 positive single stranded viral mRNA and the modified gs5* that includes Cy28 and the SV5-tag. The nucleotide sequence is shown below; red arrowhead indicates Csy4 cleavage site. The minimal 16 nt sequence specific for Csy4 binding is highlighted in light blue. B-C) Representative electrophoretic pattern (B) and quantification analysis (C) of dsRNA genome segments of rRV-wt or rRV-gs5* derived from the indicated cells at 16 hpi. Red arrow indicates gs5*, while red asterisks indicate the edited gs5*. The number of each genomic segment is indicated on the left. D) Nucleotide sequence of the gs5* edited versions gs5*-Δ27 and gs5*-Δ42. E) Relative number of gs5*-Δ27 and gs5*-Δ42 sequences in rRV-gs5*-infected MA-NSP5-Csy4 or MA-NSP2-Csy4 cells. Deleted edited forms of gs5* were RT/PCR amplified from dsRNA, cloned and sequenced (45 clones and 30 clones for MA-NSP5-Csy4 or MA-NSP2-Csy4 cells respectively). F) RT/PCR amplification of gs5*-Δ27 (174 bp) and gs5*-Δ42 (159 bp) of total RNA derived from MA-NSP5-Csy4 or MA-NSP5-Csy4-H29A cells infected with rRV-gs5* at 7 hpi. PCR was performed with primers specific for each gs5* edited version. A portion of gs11 was amplified as control.

Rotavirus genome replication in viroplasms is performed by the RNA dependent RNA polymerase (RdRp) VP1, which synthesises dsRNA using the single stranded RNA (ssRNA) of positive polarity as a template (Lu et al., 2008). Several *in vitro* evidences suggest that the terminal 3’ consensus sequence present in the viral mRNAs (5’-UGUGACC-3’) is fundamental for priming replication of genome segments (Lu et al., 2008; McDonald and Patton, 2011).

rRV-gs5* was designed to investigate the fate of virus replication and production of the mature gs5* dsRNA upon cleavage of the viral positive ssRNA by Csy4 chimeras localised to viroplasms. Analysis of the newly produced dsRNAs at 16 hpi, following rRV-gs5* infection of MA-NSP5-Csy4 cells, showed normal migration of all genome segments with the exception of gs5*, which migrated mainly as a shorter segment (80% of all gs5* band) (Figure 3B, lane 4; Figure 3C). Further passages in MA-NSP5-Csy4 cells resulted in progressive reduction of the non-edited gs5* up to complete gs5* editing after the second passage (Figure S3A,B). In contrast, infection of the parental MA104 or MA-NSP5-Csy4-H29A did not show any difference on the dsRNA migration patterns (Figure 3B, lanes 2, 3; Figure 3C). A similar gs5* deletion was observed in MA-NSP2-Csy4 cells, despite it represented only 20% of all gs5* (Figure 3B lane 5; Figure 3C), consistent with the lower nuclease activity compared to MA-NSP5-Csy4 cells (Figure 2). Genome editing of gs5* dsRNA required Csy4 localisation to viroplasms, as infection with rRV-gs5* of MA104 cells expressing the similarly active SV5-Csy4 nuclease (MA-Csy4) (Figure 2), which is not targeted to viroplasms, did not affect the migration pattern of the newly made dsRNA (Figure 3B lane 6; Figure 3C). These data suggest that RV genome editing requires cleavage of the replication intermediates present within viroplasms (Silvestri et al., 2004). The finding of the edited gs5* is remarkable, not only because it shows that nuclease mediated cleavage of genome replication intermediates can be repaired, but also because they do so with an expectedly high efficiency. Furthermore, spontaneous genome modifications in RV have been mainly described as partial duplications of the viral segments (Desselberger, 1996; Giambiagi et al., 1994; Gonzalez et al., 1989; Mattion et al., 1988; Méndez et al., 1992; Schnepf et al., 2008; Troupin et al., 2011). For both NSP5-Csy4 and NSP2-Csy4 cells, sequence analysis of the newly-edited gs5* revealed that it consisted of a mixture of two segments carrying a deletion of either 42 (gs5*-Δ42) or 27 (gs5*-Δ27) nucleotides compared to gs5* (Figure 3D), in a ratio of 55 to 45% (NSP5-Csy4) and 60 to 40% (NSP2-Csy4), respectively (Figure 3E). These data were further confirmed by transcript-specific RT/PCR detection of the two edited Δ42 and Δ27 gs5* RNAs at 7 hpi in MA-NSP5-Csy4-infected cells and not in cells expressing the nuclease inactive NSP5-Csy4-H29A protein (Figure 3F, left panel). Control gs11 was equally amplified in both infected cell lines (Figure 3F, right panel). Finding the same two gs5* deletions of 42 and 27 nucleotides with both Csy4 fusions indicated that the choice of the viral shuttle proteins (NSP2 or NSP5) does not influence the outcome of RV gs5* editing. In addition, we observed that the 3’ end of both deletions was always compatible with the Csy4 cleavage site (Figure 3D), consistent with the requirement of Csy4 nuclease activity. In both cases a G was present immediately upstream of the deletions. The two newly edited rRVs gs5*-Δ42 and gs5*-Δ27 were independently packaged into newly made viral particles and were not further mutated after 6 consecutive passages in MA104 cells (Figure S3C lanes 4, 5), as confirmed also by sequencing (Figure S3E). Consistently with the reported high specificity of Csy4 nuclease (Haurwitz et al., 2012, 2010; Sternberg et al., 2012; Young et al., 2013) *i)* deletions were observed exclusively in the RV genomic segment containing the Csy4 target sequence (Figure 3B, Figure S3A,C); *ii)* the wild type virus (rRV-wt) equally replicated in MA-NSP5-Csy4 and MA-NSP5-Csy4-H29A cells without any modification in the genome segments pattern, indicating no evident Csy4 off-target activity to the viral genome (Figure S3D).

We then asked whether the observed editing events were dependent on the position of the nuclease target sequence. In the case of the above described gs5*, Cy28 was located 95 nucleotides upstream of the 3’ end of the positive ssRNA. We thus generated a rRV carrying a similar gs5* with the same sequence containing Cy28 located 284 nucleotides upstream of the 3’ end (termed, rRV-gs5*/284) (Figure S4A). As NSP1 is dispensable for viral replication in RV-infected cells (Kanai et al., 2017; Komoto et al., 2018), the inserted modification, which generates a truncated NSP1 ORF (47 kDa), did not affect rescue of the recombinant virus.

The dsRNA profile and sequence analysis showed that almost all (95%) of gs5*/284 was edited to deleted variants when infecting MA-NSP5-Csy4 cells, but not when infecting parental MA104 cells or cells expressing the H29A Csy4 fusion (Figure S4B,C). The edited gs5*/284 segments contained, with a similar frequency, the same 42 and 27 nucleotides deletions upstream of the Csy4 cleavage site observed for gs5* (Figure S4D,E), ruling out that the position of Cy28 within the RV genomic segment affects the deletion outcome.

### CRISPR-Csy4 editing of RV gs7 and gs10

In order to investigate whether similar editing events could take place in other genome segments, two additional rRVs were obtained (Figure 4A). In the first one (rRV-gs7*), gs7 was modified inserting a C-terminal SV5-tag to NSP3 followed by the Cy28 sequence after the UGA STOP codon of the NSP3-SV5 encoded protein (gs7*; Figure 4A, upper panel). When infecting MA-NSP5-Csy4 cells, 80% of gs7* was edited, whereas no modifications were observed in other RV genomic segments or in cells expressing NSP5-Csy4-H29A (Figure 4B,C). The presence of the same deletions of 27 and 42 nucleotides and at similar proportion (38% and 62%, respectively) (Figure 4D), as observed in gs5* and gs5*/284, suggests that the editing outcome was dependent of the inserted sequence, which was identical in all three recombinant genomic segments (Figure 4A, 3A, Figure S4A).

**Figure 4.**
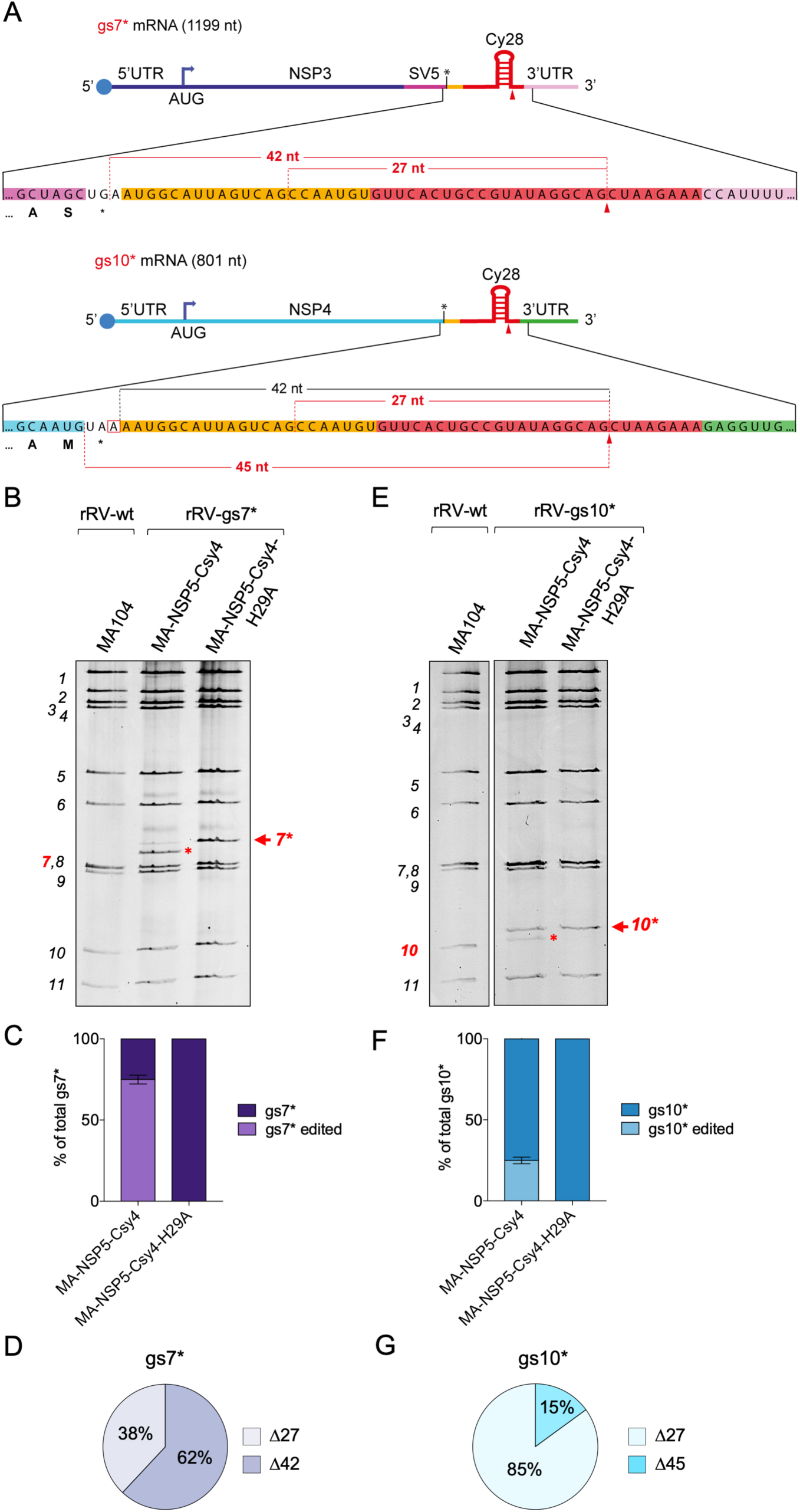
Csy4-mediated editing of rRV-gs7* and rRV-gs10*. A) Scheme of the modified gs7* (upper panel) and gs10* (lower panel). Detected deletions are indicated in red. B) Representative dsRNA electropherotypes of rRV-gs7* infecting the indicated cell lines at 16 hpi. C) Quantification analysis of the non-edited and edited gs7* present in the dsRNA electropherotypes shown in (B); data are expressed as means +/- SEM (n=3). D) relative proportion of each edited rRV-gs7* sequence. E), F) and G) are as in B), C) and D) in cells infected with rRV-gs10*. In B) and E) the number of the genomic segments are indicated on the left. The red arrows indicate gs7* and gs10*, while red asterisks indicate the respective edited versions. rRV-wt is provided as reference.

The second rRV strain contained a modified gs10 (rRV-gs10*) that encodes for NSP4 (Figure 4A, lower panel). In this case, the SV5-tag was not introduced, but the inserted sequence still contained Cy28 after the natural NSP4 UAA STOP codon (Figure 4A, lower panel). Because of the different STOP codon usage, in this design the G previously involved in the formation of Δ42 in gs5*, gs5*/284 and gs7* was not present (Figure 4A, lower panel). Upon infection of MA-NSP5-Csy4 cells, but not MA-NSP5-Csy4-H29A cells, a band of edited gs10* was detected and represented 20% of total gs10* (Figure 4E,F). For gs10* we observed the same Δ27 editing event (85%) and a new deletion of 45 nt (Δ45) (15%), which also involved a G (Figure 4G). The absence of the Δ42 editing and the appearance of the Δ45 suggest that the G nucleotide upstream of the deletion plays an important role in determining the editing outcome.

To test whether Csy4 approach was suitable also for multiplexed RV genome editing, we generated the rRV strain rRV-gs(5*-7*-10*), having the Cy28 sequence in three different RV genome segments (gs5*, gs7* and gs10*). Upon infection of MA-NSP5-Csy4 cells, but not in cells expressing the H29A fusion, rRV-gs(5*-7*-10*) was edited in all the three genome segments in a similar proportion and with the same deletions as in viruses in which only a single segment was targeted (Figure 5A, B; Figure S5A-B); as expected migration of other segments was unaffected. Taken together, these data indicate that the Csy4-targeted cleavage of RV positive ssRNA templates within viroplasms can be exploited to generate simultaneous defined editing events in multiple genome segments, in a single replication cycle.

**Figure 5.**
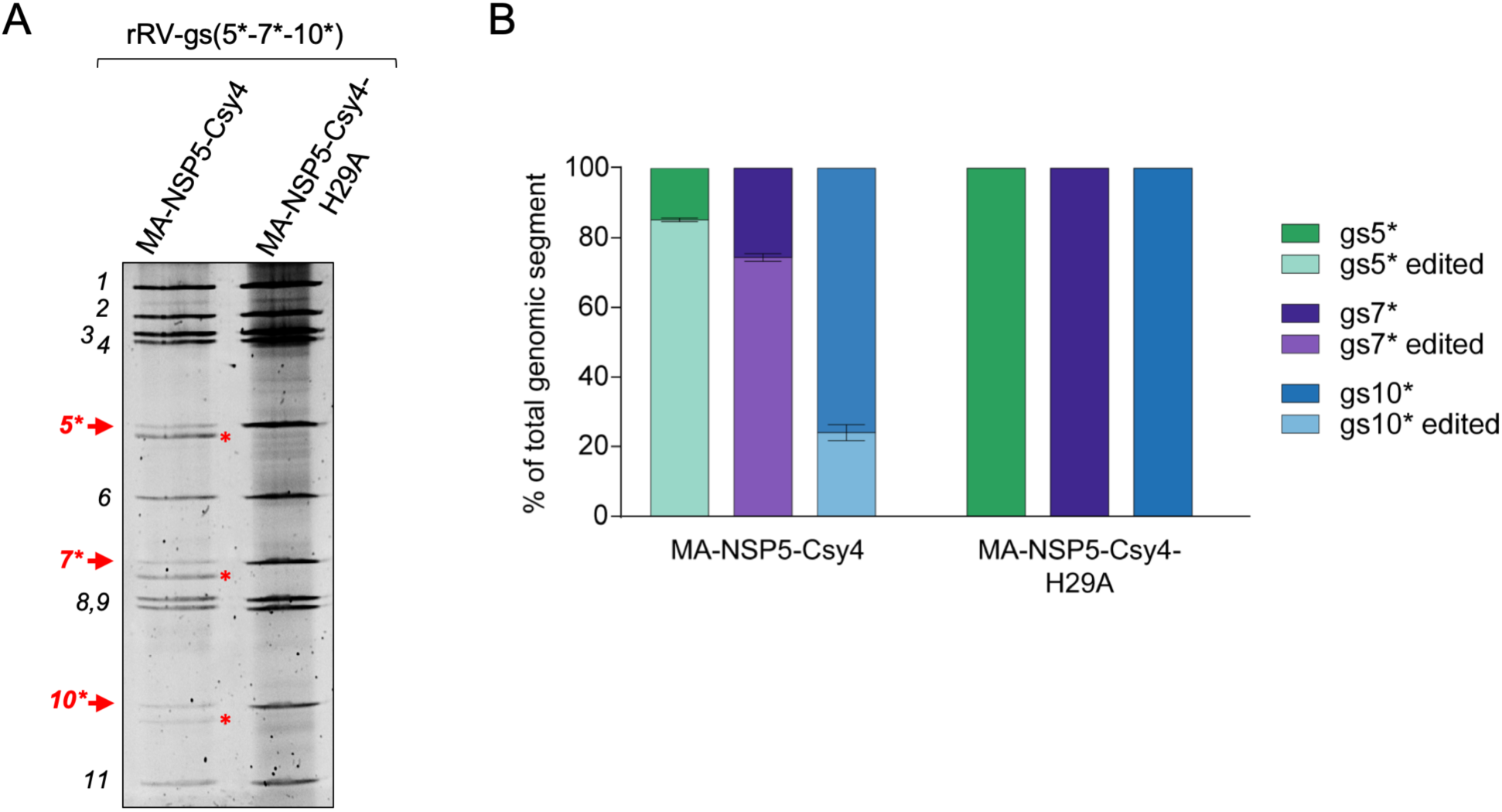
Multiplexed Csy4-mediated editing of gs5*, gs7* and gs10*. A) Representative dsRNA electropherotypes of rRV-gs(5*-7*-10*) infecting the indicated cell lines at 16 hpi. The number of genomic segments is indicated on the left. Red arrows indicate gs5*, gs7* and gs10*, while red asterisks indicate the respective edited versions. B) Quantification analysis of the non-edited and edited gs5*, gs7* and gs10*, present in the dsRNA electropherotypes shown in (A); data are expressed as means +/- SEM (n=2).

### Csy4-mediated editing to study RV secondary transcription

A hallmark of RV biology is the transcriptional activity of intermediate viral particles (Desselberger, 2014; Estes and Greenberg, 2013). Mature RVs are Triple Layered Particles (TLPs), which after entry into cells are uncoated to become transcriptionally active Double Layered Particles (DLPs). DLPs generate transcripts that serve as templates for the synthesis of dsRNA and as mRNAs to produce the whole set of viral proteins and promote the formation of viroplasms. During virus replication newly-made DLPs are assembled within viroplasms of infected cells and are supposed to start a new wave of transcription, called ‘secondary transcription’, before budding into the ER for final maturation into TLPs (Estes and Greenberg, 2013).

Beyond some indirect evidences, to what degree secondary transcription by the newly-made DLPs and the consequent translation contribute to the overall production of viral proteins remains a largely unknown aspect in RV replication (López et al., 2005). Csy4-mediated editing of the RV genome provides a novel approach to study secondary transcription, as it allows monitoring expression of the edited genome segment in a single replication cycle. In fact, upon transcription the newly generated edited segment produces an mRNA different from the one of the original infective particles.

To detect proteins produced exclusively by the edited mRNA we engineered a new rRV (herein named rRV-gs5*-HA), where the gs5*-HA contained a slightly shorter Csy4 target sequence (Cy23) and the HA-tag coding sequence downstream of the NSP1-SV5 STOP codon (Figure 6A). Cy23 contains the essential 16 nt hairpin that is recognised and efficiently cleaved by Csy4 (Sternberg et al., 2012). gs5*-HA was designed to express HA-tagged isoforms of NSP1-SV5 after in-frame deletions of at least 27 nucleotides, which can eliminate the STOP codon (Figure 6A). Upon infection of MA-NSP5-Csy4 cells two different editing events of gs5*-HA, corresponding to deletions of 36 and 21 nucleotides were detected, with a total editing efficiency of 80% (Figure 6B, C; Figure S6). The gs5*-HA Δ36 transcript was the most frequent (85%) (Figure 6D) and started to be detected at an early time post-infection (5 hpi), increasing up to be preponderant (63%) compared to the non-edited transcript at later time points (6-9 hpi) (Figure 6E). The Δ36 editing was expected to result in a 61.4 kDa HA-tagged NSP1-SV5 protein, while the Δ21 and the non-edited gs5*-HA encoded a 60.4 kDa NSP1-SV5 isoform (Figure 6A). Consistently, western blot analysis detected a single band of HA-tagged NSP1-SV5 in MA-NSP5-Csy4 cells from 7 hpi (Figure 6F), but not in cells expressing the inactive NSP5-Csy4-H29A, where gs5*-HA was not edited (Figure 6F). As only newly made particles can contain an edited gs5*-HA, expression of NSP1-SV5-HA allowed monitoring translation of the newly produced transcripts derived from secondary transcription. Time course analysis using anti-SV5 antibody which labels both HA-edited and non-edited NSP1 showed that from 5 to 9 hpi, production of NSP1-SV5-HA progressively increases reaching almost 50% of the total NSP1 at 7 hpi and become predominant at later time points (9 hpi), contributing for more than 70% of the whole production of the viral protein (Figure 6G,H). Of note, the contribution of the secondary transcription and derived translation is likely underestimated because of the incomplete editing of gs5*-HA and the generation of the Δ21 isoform (15%), which does not remove the stop codon after SV5-tag, thus preventing expression of the longer HA-tagged protein.

**Figure 6.**
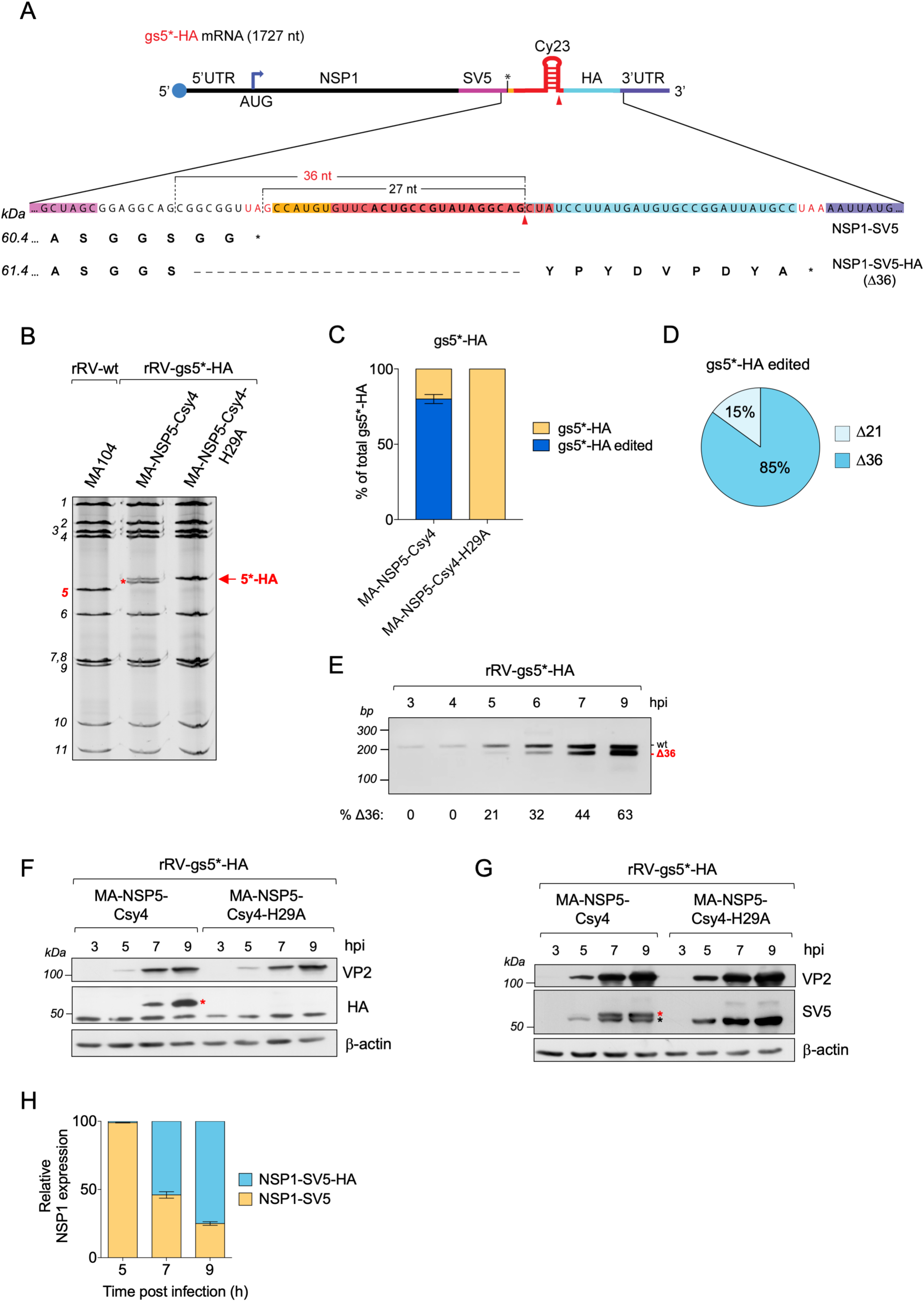
Efficient Csy4-mediated detection of secondary transcription. A) Scheme of the 1727 nt long mRNA of gs5*-HA, showing the potential minimum in-frame deletion (Δ27) to express the HA-tagged NSP1 and the very frequently detected Δ36 (red) editing product. The C-terminus amino acid sequence and the expected molecular mass of NSP1-SV5 and NSP1-SV5-HA proteins are shown. Arrowhead indicates the Csy4 cleavage site. B) Representative electrophoretic pattern of dsRNA genome segments of the rRV-gs5*-HA strain replicated in the indicated cell lines at 15 hpi. Arrow and red asterisk label, respectively, the non-edited and edited gs5*-HA. C) Quantification analysis of gs5*-HA from the electrophoretic patterns shown in (B). Data are expressed as means +/- SEM (n=3). D) Frequency of gs5*-HA-Δ21 and gs5*-HA-Δ36 sequences (36 clones) generated in rRV-gs5*-HA-infected MA-NSP5-Csy4 cells at 9 hpi. E) Representative transcript specific RT/PCR amplification (see Materials and Methods) of gs5*-HA (213 bp) and gs5*-HA-Δ36 (177 bp) derived from MA-NSP5-Csy4 cells infected with rRV-gs5*-HA at the indicated times post infection. The percentage of Δ36 transcripts represent the mean of n=3 experiments. F) Western blot of the time course expression of NSP1-SV5-HA (detected with anti-HA) from extracts of MA-NSP5-Csy4 and MA-NSP5-Csy4-H29A cells infected with rRV-gs5*-HA (middle panel). Expression of the control viral protein VP2 is shown (upper panel). β-actin was used as loading control. G) Western blot of the same extracts used in (F) developed with anti-SV5 (middle panel). Red and black asterisks indicate the HA-tagged and non-tagged NSP1-SV5, respectively. H) Quantification of the relative expression of the two NSP1 isoforms shown in (G). Data are expressed as means +/- SEM (n=3).

Encouraged by the positive result with the rRV gs5*-HA, we produced a new rRV with a gs5 modified to encode EGFP after the HA-tag (rRV-gs5*-HA-EGFP*) (Figure 7A). Also in this case, EGFP would be expressed only upon Csy4-mediated in-frame deletion of at least 27 nucleotides. In addition, compared to gs5*-HA, we modified the sequence upstream Cy23 removing two G nucleotides, including the one previously found involved in the formation of the Δ21 and Δ27 deletions. The size of gs5*-HA-EGFP* was too large to clearly detect the deleted edited segment in the dsRNA gel migration profile (Figure 7B). However, cytofluorimetric analysis (Figure 7C) and confocal fluorescence images (Figure 7D,E) showed 15% of EGFP positive cells in rRV-infected MA-NSP5-Csy4 cells at 12 hpi. Analysis by Sanger sequencing decomposition (Brinkman et al., 2014) indicates that the only significant editing event was the Δ36 deletion (Figure S7A). As expected, EGFP fluorescence was detected only in cells sustaining viral replication (anti-NSP2) (Figure 7D) and was strongly compromised upon inhibition of viroplasms formation by treatment with the proteasome inhibitor MG132 (Contin et al., 2011) (Figure 7E; Figure S7C,E). On the other hand, EGFP positive cells were absent in control infected MA-NSP5-Csy4-H29A cells (Figure 7C,D; Figure S7C,F). We followed the kinetics of NSP1-EGFP produced by secondary transcription in virus-infected cells by live-cell imaging (Figure 7E,F, Figure S7B,C,D and Supplement Movie 1). EGFP fluorescence was detected as early as 6 hpi and increased over time reaching a plateau in the number of EGFP positive cells from 13 hpi (Figure 7E,F and Supplement Movie 1). Our data thus indicate that secondary transcription plays an important role in viral proteins production and demonstrate that Csy4-mediated RV genome editing can be used to study this process by live imaging.

**Figure 7.**
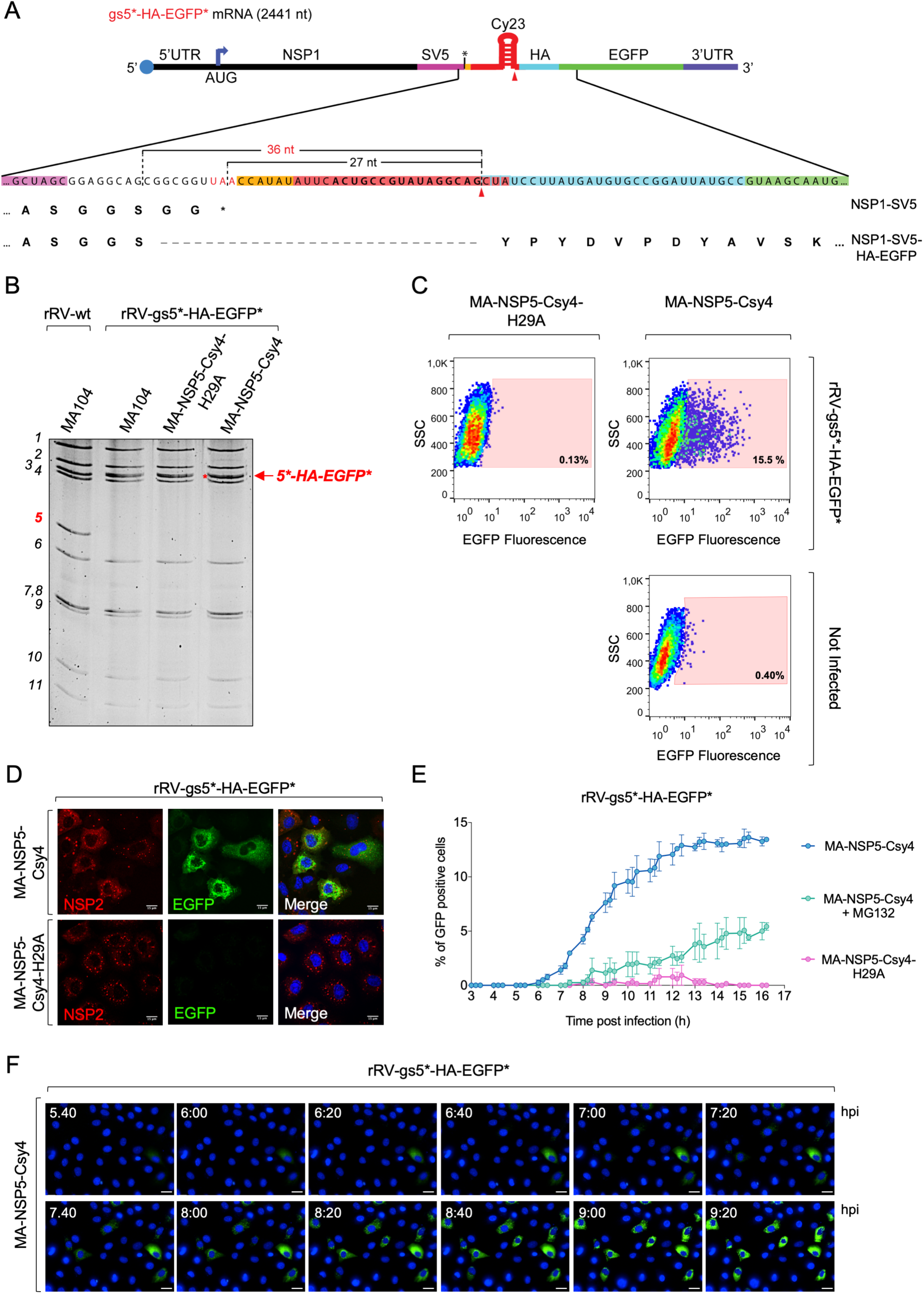
Live-cell imaging of Csy4-mediated RV secondary transcription and translation. A) Scheme of the 2441 nt long mRNA of gs5*-HA-EGFP* showing the minimal in-frame Δ27 and expected Δ36 deletion required for EGFP expression. B) Representative dsRNA electropherotype of rRV-gs5-HA-EGFP* strain infecting the indicated cell lines at 16 hpi. The number of genomic segments is indicated on the left. The red arrow indicates gs5*-HA-EGFP*. rRV-wt is provided as reference. C) Cytofluorimetric analysis of MA-NSP5-Csy4-H29A and MA-NSP5-Csy4 cells infected with rRV-gs5*-HA-EGFP* (MOI 5) at 12 hpi. The percentage of EGFP positive cells is indicated. D) Immunofluorescence of MA-NSP5-Csy4 and MA-NSP5-Csy4-H29A cells infected with rRV-gs5*-HA-EGFP* at 12 hpi. Infected cells and viroplasms were detected with anti-NSP2 (red) and nuclei with DAPI (blue). As expected, NSP1-EGFP (green) does not localise to viroplasms. Scale bar 13 μm. E) Percentage of EGFP fluorescent cells (all cells scoring a number of NSP1-EGFP positive spots ≥ 1) over time (from 1 to 17 hpi) of MA-NSP5-Csy4 (non-treated and treated with MG132) and MA-NSP5-Csy4-H29A cells infected with rRV-gs5-HA-EGFP* (MOI 5) monitored by live-cell imaging. F) Time course of EGFP fluorescence single field micrographs (from 5:40 to 9:20 hpi) of the data shown in E). One field (50 cells) where EGFP fluorescence was reconstituted is shown. DAPI stained nuclei shown in blue. Scale bar 50 μm.

## DISCUSSION

CRISPR-Cas nucleases, especially *Streptococcus pyogenes* CRISPR-Cas9, have become the tool of choice to edit genomes of cells and organisms (Makarova et al., 2018; Song et al., 2016; Zhang et al., 2018). Consistently with their natural role, CRISPR-nucleases have also been exploited to disrupt and manipulate viral genomes with various purposes. DNA viruses including Hepatitis B virus (HBV), Herpesviruses (HSV), Human Papilloma Virus (HPV), as well as lentivirus (HIV-1) and retroviruses have been efficiently targeted by CRISPR-Cas in different cell-culture and *in vivo* systems (de Buhr and Lebbink, 2018; Chen et al., 2018; van Diemen et al., 2016; Lao et al., 2018; Li et al., 2018; Schiwon et al., 2018; Xiao et al., 2019; Yoshiba et al., 2019). In these cases, the edited target molecule is the viral DNA genome or its DNA replication intermediate (Goetze et al., 2017; Wang et al., 2016b, 2016a).

Members of the type VI CRISPR-Cas13 family of endoribonuclease have been used to specifically target and degrade viral single-stranded RNAs of HIV-1 and HCV, interfering with viral infection (Cox et al., 2017; Guo et al., 2015). Similarly, CRISPR-Csy4 was used to block infection of a model HIV-1 virus (Guo et al., 2015). However, there are no reports showing the use of CRISPR-nucleases or other homing nuclease in targeting and editing a dsRNA genome. Indeed, the peculiar replication mechanism of RV in hardly accessible viral factories poses several challenges for the effective deployment of any nucleic acid editing tool (Silvestri et al., 2004). In addition, there are no known repair mechanisms for cleaved RV genome, nor sequence specific programmable CRISPR nucleases targeted to dsRNA published yet. We paved the way for nuclease-mediated genome editing of dsRNA viruses using the small and highly specific CRISPR-Cas type I RNA endonuclease Csy4 of *P. Aeruginosa* (Sternberg et al., 2012) to cleave the positive ssRNA intermediates formed during RV dsRNA genome replication (Borodavka et al., 2017).

To successfully target the viral genome, we had to delivered Csy4 to viroplasms, the intracellular compartment of RV replication, where the positive ssRNA replication intermediates are located (Desselberger, 2014; Estes and Greenberg, 2013; Patton et al., 2006). This was obtained by fusing Csy4 to the viroplasm-localising viral proteins NSP5 and NSP2 (Eichwald et al., 2004, 2012; Fabbretti et al., 1999). Among the 11 RV genomic segments we selected three of them, gs5, gs7 and gs10, to be engineered to contain the Csy4 target sequence in the genomic RNA plus strand, positioning it in different locations within the genomic segments. The viroplasm-localised nuclease Csy4 produced discrete deletions ranging from 21 to 45 nt with efficiency up to 95% in a single round of infection, which increased up to 100% in following rounds of infection in NSP5-Csy4 expressing cells.

Remarkably, a virus carrying three segments (gs5, gs7 and gs10) with the Csy4 target sequence was also very efficiently edited upon Csy4-mediated cleavage in all the three segments, thus offering the opportunity to specifically and simultaneously induce *in vivo* editing isoforms in different viral genes. Editing was detected exclusively on genome segments carrying the stem-loop target sequence and not on the other segments, thus indicating that Csy4 fused to either NSP5 or NSP2 can be safely used in the context of RV infection as no off-targeting cleavage of the wild type virus genome was observed.

The presence of editing events exclusively when Csy4 was localised to viroplasm indicates that the Csy4-mediated editing of the dsRNA segment takes place in the RV viral factories and suggest an involvement of the double stranded RNA replication step. Consistently, RNA interference against RV positive ssRNAs have never been reported to cause deletions within the target genome segments, probably because viroplasm-localised positive ssRNAs intermediates are not accessible to the RISC complex (Silvestri et al., 2004).

RV replication requires the polymerase activity of the RdRp VP1, together with the core protein VP2 and the positive ssRNA as template (Ding et al., 2019; Lu et al., 2008; Patton et al., 1997). The exact organisation on how RV dsRNA replication occurs is still debated (Borodavka et al., 2018; Ding et al., 2019; Gallegos and Patton, 1989), but two different models have been proposed: i) pre-assembled cores containing VP1, VP3 and VP2 allow recruitment of single copies of positive ssRNAs of each segment, which are then converted into dsRNA following synthesis of the negative strands (Long et al., 2017) or, ii) positive ssRNA replicates in pre-cores intermediates containing VP1-VP3 complexes associated to VP2 decamers, but lacking a complete VP2 shell (Gallegos and Patton, 1989; McDonald and Patton, 2011). Our data favour this second scenario, as the positive ssRNAs appear to be available for cleavage by NSP2- or NSP5-Csy4 fusions within viroplasms during the initial steps of RV genome replication.

All the edited dsRNA genome segments preserved the sequence downstream of the Csy4 cleavage site, restoring the 3’UTR. This represents the first evidence of a repair mechanism for cleaved viral ssRNA of a virus with a dsRNA genome. In all cases, the edited sequences contained a deletion where the 3’ end can always be aligned to start exactly from the Csy4 cleavage site, after the G positioned at the base of the stem of the Csy4 RNA target structure. In addition, all deletions in the targeted genome segments initiate after a G nucleotide and, in some cases, have a micro-homology (3-6 nt) to the last nucleotides of the Csy4 binding site (Figure 8). The G nucleotide seems to be essential for this efficient viral RNA repair as removing a specific G involved in a defined deletion (Δ42 in gs10*) completely prevented that editing outcome and enabled a deletion involving another proximal G (Δ45 in gs10*). Similarly, replacing the G involved in the Δ27 editing of gs5*-HA (G to A) led to the generation of new editing events (Δ36 and Δ21), further suggesting the essential role of the G nucleotide.

**Figure 8.**
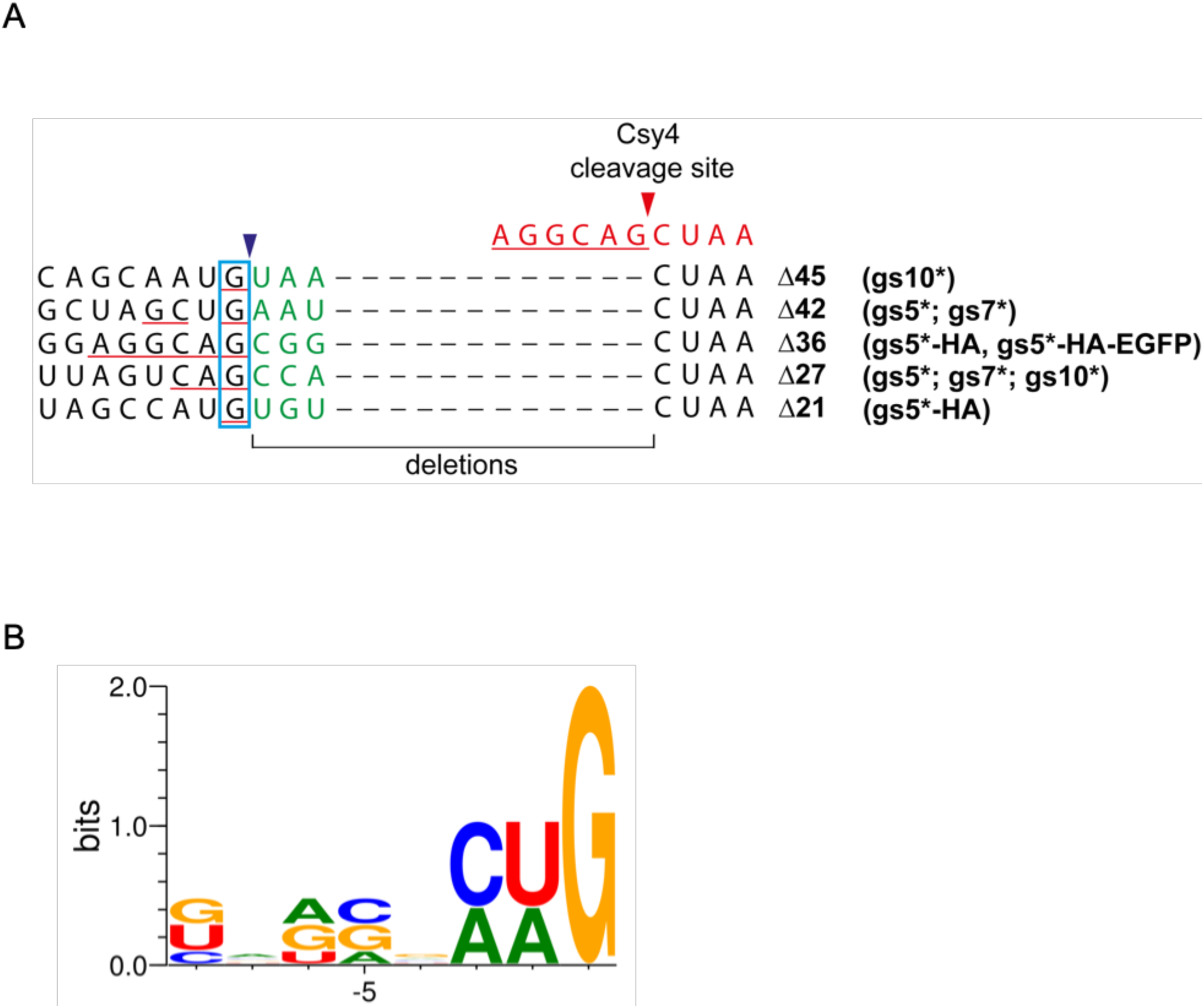
Schematic representation of the Csy4-mediated deletions found in edited RV genome segments. A) Schematic RNA sequences of the edited RV genome segments. The 5’ ends of the deletions are shown in green. The conserved G immediately upstream of the deletions is highlighted in blue. Nucleotides with microhomology to the Csy4 cleavage site are underlined in red. Red and dark blue arrows indicate deletion boundaries. Sequence of the Csy4 3’ cleavage site is shown in red. B) Frequency of the first 8 nucleotide upstream of the deletions shown in A), was calculated and plotted as a WebLogo3 (Crooks et al.).

It is therefore possible that the repair of viral RNA forming these deletions is the consequence of a second upstream cleavage by an unidentified nuclease that is followed by a cytosolic splicing ligation of the 5’ and 3’ positive ssRNA fragments, or alternately, an unusual and so-far undescribed VP1-mediated repair, both of which may take place within viroplasms during genome replication. Regardless of which is the exact mechanism, the presence of G nucleotides appears to dictate the editing events.

We applied the *in vivo* editing of the dsRNA RV genome to address an important question related to the RV replication cycle, such as the contribution of transcription by the newly-made viral DLPs during infection (Desselberger, 2014; Estes and Greenberg, 2013; López et al., 2005), which cannot be addressed only by the reverse genetics available protocols. Several RNA viruses show secondary transcription as a consequence of transcription of subgenomic RNAs at different times during the replication cycle (Repik et al., 1974; Skehel and Hay, 1978). In the case of RV and of other members of the *Orthoreoviruses* this question is not easily addressed as the progeny virus has exactly the same composition as the infecting particles (Acs et al., 1971; Sakuma and Watanabe, 1971; Skup and Millward, 1980; Zweerink and Joklik, 1970).

We thus built a rRV containing a gs5* encoding an EGFP reporter, which could be expressed only following Csy4-mediated editing. We successfully visualised the onset of the products of secondary transcription by live-cell imaging in a large proportion of infected cells.

Thanks to Csy4-mediated editing of the rRVs gs5*-HA and gs5*-HA-EGFP* we detected secondary transcription made by newly made DLPs as soon as 5 hpi, while the translation products of secondary transcripts were clearly visible slightly later (6-7 hpi). Using the rRV gs5*-HA as an example to monitor products coming from primary and secondary transcription, we could show that the contribution of secondary transcription and translation at 9 hpi is responsible for the large fraction of newly made gs5 derived transcripts (63%) and proteins (70%) produced. These results suggest that the efficiency of virus replication is affected by the capacity of the newly assembled DLPs to produce mRNAs within viroplasms that are then efficiently exported for translation. In fact, MG132-mediated impairment of viral replication, by inhibiting viroplasm formation, strongly compromises the detection of products of secondary transcription, thus suggesting that these editing events require active viral replication.

The recent development of different reverse genetics protocols for RV represents a powerful tool to investigate several aspects of the virus biology and replication (Kanai et al., 2017; Komoto et al., 2018). We believe that nuclease-mediated editing of dsRNA virus could, at the same time, overcome current limitations of reverse genetics systems or be combined with them for several new possible applications, such as producing many edited virus variants unachievable directly by reverse genetics.

Overall, our data represent the first description of a nuclease-mediated editing of a dsRNA viral genome. The *in vivo* generation of a new progeny of recombinant RVs with edited genomes, paves the way for the harnessing of this tool to study different aspects of viral replication, as well as its use for the identification of therapeutic drugs targeting assembly of DLPs or pathways directly associated to secondary transcription.

## MATERIALS AND METHODS

### Cell lines

MA104 (embryonic African green monkey kidney cells ATCC CRL-2378.1) cells were cultured in Dulbecco’s Modified Eagle’s Medium (DMEM) (Life Technologies) supplemented with 10% Fetal Bovine Serum (FBS) (Life Technologies) and 50 μg/ml gentamycin (Biochrom AG).

MA104-NSP5-Csy4 (MA-NSP5-Csy4), MA104-NSP5-Csy4-H29A (MA-NSP5-Csy4-H29A), MA104-Csy4 (MA-Csy4), MA104-Csy4-H29A (MA-Csy4-H29A), MA104-NSP2-Csy4 (MA-NSP2-Csy4), MA104-NSP2-mCherry (MA-NSP2-mCherry) stable transfectant cell lines were grown in DMEM containing 10% FBS, 50 μg/ml gentamycin and 5 μg/ml puromycin (Sigma-Aldrich).

BHK-T7 cells (<u>B</u>aby <u>H</u>amster <u>K</u>idney stably expressing <u>T7</u> RNA polymerase) were cultured in Glasgow medium supplemented with 5% FBS, 10% Tryptose Phosphate Broth (TPB) (Sigma-Aldrich), 50 μg/ml gentamycin, 2% Non-Essential Amino Acids (NEAA) and 1% Glutamine.

### Plasmid construction

RV plasmids pT7-VP1-SA11, pT7-VP2-SA11, pT7-VP3-SA11, pT7-VP4-SA11, pT7-VP6-SA11, pT7-VP7-SA11, pT7-NSP1-SA11, pT7-NSP2-SA11, pT7-NSP3-SA11, pT7-NSP4-SA11, and pT7-NSP5-SA11 (Kanai et al., 2017; Komoto et al., 2018) were used to rescue recombinant RVs by reverse genetics.

pPB-NSP5-SV5-Csy4 and pPB-SV5-Csy4 plasmids were obtained from a GenParts DNA fragment (Genscript) containing NSP5-SV5-Csy4 and SV5-Csy4 and inserted in the pPB-MCS vector (VectorBuilder) using BamHI-XmaI restriction enzymes sites.

pPB-NSP5-SV5-Csy4-H29A and pPB-SV5-Csy4-H29A plasmids carrying two nucleotides substitutions C82G and A83C in the Csy4 ORF were generated by QuikChange II Site-Directed Mutagenesis (Agilent Technologies) from the pPB-NSP5-SV5-Csy4 and pPB-SV5-Csy4 respectively using Csy4-H29A-FOR and Csy4-H29A-REV primers (Figure Supplement Data 1).

A GenParts DNA fragment containing the NSP2-SV5-Csy4 was inserted into pPB-MCS vector using NotI-EcoRI restriction enzymes sites to harbour the pPB-NSP2-SV5-Csy4.

pEGFP-Cy28 was generated inserting Cy28 after the ATG into the pEGFP-C1 plasmid (Addgene) using the pEGFP-Cy28-FOR and pEGFP-Cy28-REV primers (Figure Supplement Data 1) for QuikChange II Site-Directed Mutagenesis.

pT7-gs5* was generated inserting a GenParts DNA fragment into the pT7-NSP1-SA11 using PacI-BamHI restriction enzymes sites. This GenParts fragment contains the SV5 tag and Cy28 at position 1518 of the gs5.

pT7-gs7* was generated inserting a GenParts DNA fragment into the pT7-NSP3-SA11 using SnaBI-SacI restriction enzymes sites. The GenParts fragment contains the SV5 tag and Cy28 at position 970 of the gs7.

pT7-gs10* was modified to insert Cy28 sequence using gs10*-Cy28-FOR and gs10*-Cy28-REV primers by QuikChange II Site-Directed Mutagenesis.

For the generation of pT7-gs5*/284, a GenParts DNA fragment containing the SV5 tag and Cy28 was inserted into pT7-NSP1-SA11 using MfeI-BamHI restriction enzymes sites.

pT7-gs5*-HA was obtained inserting a GenParts DNA fragment including the SV5 tag, Cy23 and the HA tag in plasmid pT7-NSP1-SA11 using MfeI-BamHI restriction enzymes sites.

pT7-gs5*-HA-EGFP* was generated cloning a GenParts DNA fragment into the pT7-NSP1-SA11 using MfeI-BamHI. The GenParts contains the SV5 tag, Cy23, the HA tag and the EGFP ORF upstream of the 3’UTR of gs7 and HDV ribozyme.

### Generation of stable cell lines

MA-NSP2-mCherry was previously described (Papa et al., 2019). MA-NSP5-Csy4, MA-NSP5-Csy4-H29A, MA-Csy4, MA-Csy4-H29A, MA-NSP2-Csy4 were similarly generated using the PiggyBac Technology (Papa et al., 2019; Yusa et al., 2011). Briefly, MA104 cells (10^5^) were seeded in a 12 Multi-well plate, transfected the next day with the pCMV-HyPBase (0.5 μg) and the respective transposon plasmids: pPB-NSP5-SV5-Csy4, pPB-NSP5-SV5-Csy4-H29A, pPB-SV5-Csy4, pPB-SV5-Csy4-H29A and pPB-NSP2-SV5-Csy4, using a ratio of 1:2.5 respectively with Lipofectamine 3000 (Sigma-Aldrich) according to the manufacturer’s instructions.

Cells were maintained in DMEM supplemented with 10% FBS for 3 days and then incubated with DMEM supplemented with 10% FBS and 10 μg/ml puromycin for 4 days for selection.

### Rescue of recombinant RVs from cloned cDNAs

To rescue recombinant RV strain SA11 (rRV-wt), monolayers of BHK-T7 cells (4 × 105) cultured in 12-well plates were co-transfected using 2.5 μL of TransIT-LT1 transfection reagent (Mirus) per microgram of DNA plasmid. Each mixture comprised 0.8 μg of SA11 rescue plasmids: pT7-VP1, pT7-VP2, pT7-VP3, pT7-VP4, pT7-VP6, pT7-VP7, pT7-NSP1, pT7-NSP3, pT7-NSP4, and 2.4 μg of pT7-NSP2 and pT7-NSP5 (Kanai et al., 2017; Komoto et al., 2018). Furthermore 0.8 μg of pcDNA3-NSP2 and 0.8 μg of pcDNA3-NSP5, encoding NSP2 and NSP5 proteins, were also co-transfected to increase rescue efficiency (Papa et al., 2019). To rescue recombinant rRVs encoding NSP1 and NSP3 mutants, pT7-gs5*, pT7-gs5*/284, pT7-gs5*-HA, pT7-gs5*-HA-EGFP*, pT7-gs7*, pT7-gs10* plasmids were used instead of pT7-NSP1-SA11, pT7-NSP3-SA11, pT7-NSP4-SA11 respectively.

Cells were co-cultured with MA104 cells for 3 days in FBS-free medium supplemented with trypsin (0.5 μg/ml) (Sigma Aldrich) and lysed by freeze-thawing. 300 μl of the lysate was transferred to fresh MA104 wt cells, further cultured at 37°C for 4 days in FBS-free DMEM supplemented with 0.5 μg/ml trypsin until a clear cytopathic effect was visible. The modified genome segments of rescued recombinant rotaviruses were sequenced.

### Replication kinetics of recombinant viruses

MA104 cells were seeded into 24-well plates, infected with rRVs at MOI of 0.5 for multi-step growth curve experiments. Cells were harvested after 6, 12, 24, 36 hours post infection and lysed by freeze-thawing three times and activated with trypsin (1 μg/ml) for 30 min at 37°C. The lysates were used to infect monolayers of MA104 stably expressing the fusion protein NSP2-mCherry (Papa et al., 2019). The MA-NSP2-mCherry cells were seeded in μ-Slide 8 Well Chamber Slide-well (iBidi GmbH) and infected with dilutions of virus-containing lysates. Cells were then fixed 5 hours after the infection for 15 min with 4% paraformaldehyde. Nuclei were then stained with ProLong Diamond Antifade Mountant with DAPI (Thermo Scientific). Samples were imaged using a confocal setup (Zeiss Airyscan equipped with a 63x, NA=1.3 objective). Each viroplasms-containing cell was counted as one focus-forming unit (FFU). The average of cells with viroplasms of six fields of view per each virus dilution was determined and the total number of cells containing viroplasms in the whole preparation was estimated. The virus titer was determined as previously described (Eichwald et al., 2012).

### Western blot

Samples from rRV-infected cells were separated by reducing 10% or 7-15% gradient SDS-PAGE, transferred to polyvinylidene difluoride (PVDF) membranes (Millipore) and blocked in a 5% milk solution in PBS (PBS-milk) for 30 minutes. SV5-tagged proteins were detected using mouse anti-SV5 mAb (1:5000, Life Technologies); HA-tagged proteins were detected by anti-HA mAb (clone HA-7, 1:5000, Sigma-Aldrich); NSP5 and VP2 proteins were detected using anti-NSP5 guinea pig serum (1:5000) or anti-NSP5 roTag mAb and anti-VP2 guinea pig serum (1:2000) (Contin et al., 2010; Eichwald et al., 2004). As secondary antibodies were used HRP-conjugated goat anti-guinea pig (1:10000; Jackson ImmunoResearch) or goat anti-mouse anti-mouse IgG goat antibodies (1:10000; KPL).

Mouse HRP-conjugated anti-GAPDH mAb (clone sc-47724, Santa Cruz Biotechnology, 1:1000) and Mouse HRP-conjugated anti-actin mAb (clone AC-15, Sigma-Aldrich, 1:40000) were used as loading controls. Membranes were developed by Enhanced ChemiLuminescence System (Pierce ECL-Western blotting system, ThermoFisher-Pierce). Quantification analysis of bands were carried out using ImageLab 6.0.1 (BioRad).

### Immunofluorescence

Immunofluorescence experiments were performed using μ-Slide 8 Well Chamber Slide-well (iBidi GmbH). Cells were washed three times with PBS and then incubated with PFA (paraformaldehyde) 3.7 % in PBS for 15 minutes, cells were then incubated with 0.1% Triton X-100 (Sigma) in PBS for 5 minutes followed by incubation with 1% PBS-BSA for 30 minutes. For the detection of proteins, cells were incubated with primary antibody diluted in 1% PBS-BSA for 1 hour, washed three times with PBS and then incubated with secondary antibody for 45 minutes. Nuclei were then stained with ProLong™ Diamond Antifade Mountant with DAPI (Thermo Scientific) and the slides were imaged using a confocal setup (Zeiss Airyscan equipped with a 63x, NA=1.3 objective). The antibodies were used with the following dilutions: mouse anti-SV5 mAb (1:1000); anti-NSP5 guinea pig serum 1:1000; anti-NSP2 guinea pig serum 1:200 (Contin et al., 2010; Eichwald et al., 2004); Alexa Fluor 488-conjugated anti-mouse, 1:500 (Life Technologies), and TRITC-conjugated anti-guinea pig, 1:500 (Life Technologies). Quantification analysis were performed using Image J software.

### Electrophoresis of viral dsRNA genomes

Cells infected with the recombinant viruses at MOI of 5 and were lysed 16 hours post infection. Total RNA was extracted with RnaZol® (Sigma-Aldrich) according the manufacturer’s protocol. The RNA was run on a 10% (wt/vol) poly-acrylamide gels (PAGE) for 2 hours at 180 Volts. The gel was than stained with ethidium bromide (1μg/ml) and the dsRNA pattern was visualised using the ChemiDoc Imaging System. Quantification analysis of bands were carried out using ImageLab 6.0.1.

### dsRNA extraction from polyacrylamide gels

The bands of interest from the run of dsRNA on a polyacrylamide gel were cut using a blade and crushed with a pipette in 1.5 ml tubes. 300 μl of a gel elution buffer (0.5 M EDTA, 0.1% SDS, 0.5 mM NH_4_OAc) was added to the RNA samples which were mixed for 2 hours to allow the RNA to exit from the polyacrylamide gel. Then, samples were centrifuged for 3 times at 5000 g for 1 minute to pellet the acrylamide residues. In order to precipitate the RNA, 1 volume of 3M NaOAc pH 5.2 and 3 volumes of 100% EtOH were added to each sample and centrifuged at 16000 rcf for 30 minutes. After 2 washes with 2 volumes of 70% EtOH in water, the RNA pellet was resuspended in an appropriate volume of RNase free water. The dsRNA was denaturated, retro-transcribed, amplified and cloned into a pcDNA 3.1 plasmid (Invitrogen) for the sequencing experiments.

### Cytofluorimetry

MA-NSP5-Csy4 and MA-NSP5-Csy4-H29A were infected with rRV-gs5*-HA-EGFP* at MOI of 5 and collected 12 hours post infection. Cells were trypsinised and washed twice with PBS followed by centrifugation for 5 minutes at 200 rcf. After centrifugation, the cells were resuspended in PBS and the EGFP fluorescence was analysed using a FACS Calibur cytofluorimeter (BD Biosciences).

### Live Cells imaging

MA-NSP5-Csy4 and MA-NSP5-Csy4-H29A cells (2×10^3^) were seeded into CellCarrier-96 Ultra Microplates and infected with rRV-gs5*-HA-EGFP* at MOI of 5. After 1 hour, cells were stained with DAPI and live-cell imaging was started. Cells were imaged at 40x magnification (Olympus 40x NA 0.95) with a PerkinElmer Operetta High content microscope under controlled environmental conditions (37°C, 5% CO_2_). Image acquisition was performed with intervals of 20 min for a total of 12 hours. Image were analyzed using Columbus analysis software (Perkinelmer) and total cell number, number of EGFP positive spots per cell and % of EGFP positive cells (number of NSP1-EGFP positive spots ≥ 1) were calculated for each time point at single cell level.

### Sequencing of edited RV genomic segments

After extraction of RNA either from cells or polyacrylamide gels, the RNA was subjected to RT-PCR and the derived-cDNAs were amplified using NSP1-FOR and NSP1-REV primers for gs5* and gs5*HA. gs10* and gs7* were amplified using gs10*-FOR, gs10*-REV, gs7*-FOR, gs7*-REV respectively (Figure Supplement Data 1). All primers contain HindIII and XhoI restriction enzyme sites at the 5’ and 3’ end respectively. The PCR products were either sequenced and analysed using the TIDE software (Brinkman et al., 2014) or gel purified before HindIII-XhoI digestion and cloning into pcDNA 3.1 plasmid.

The bacterial colonies were checked by colony PCR using primers used for cloning. The colonies-derived PCR products were gel purified and sequenced.

### PCR using splicing primers

After RNA extraction and RT-PCR, the derived cDNAs were subjected to PCR using gs5*-FOR and gs5*Δ42-REV or gs5*-FOR and gs5*Δ27-REV to specifically detect gs5*-Δ42 or gs5*-Δ27 respectively (Figure Supplement Data 1). The transcript specific PCR amplification for gs5*-HA (213 bp) and gs5*-HA-Δ36 (177 bp) from cDNA derived from MA-NSP5-Csy4 infected with rRV-gs5*-HA at different time post infection (hpi), was performed using the gs5*-HA-FOR and gs5*-HA-REV primers (Figure Supplement Data 1). While gs5*-HA-FOR anneals to all the forms of gs5*-HA transcript, the last 4 bases at the 3’ end of gs5*-HA-REV (5’-CTGC-3’) anneals only with gs5*-HA and with gs5*-HA-Δ36 but not with gs5*-HA-Δ21, allowing the specific amplification of only gs5*-HA and gs5*-HA-Δ36.

The PCR products were run on 2.5% agarose gel in TBE 1X (45 mM Tris-borate/1 mM EDTA) for 20 minutes at 120V. Quantification analysis of bands were carried out using ImageLab 6.0.1.

## Data and materials availability

Processed data associated with this study are present in the paper, in the Figure Supplement Materials or are available from the authors upon reasonable request.

## Acknowledgements

The authors are grateful to Shan Tang and Josè Luis Slon Campos for critical reading of this manuscript, and Naoto Ito for providing BHK-T7 cells. This work was supported by ICGEB intramural funding. G.Pa and E.S. were supported by ICGEB pre-doctoral fellowship.

## Author Contributions

All experiments were conducted by G.Pa. and L.V. with the exception of live cell imaging analysis (G.Pa., L.B., E.S. and M.G.). O.R.B., G.Pa. and G.Pe. planned the project and wrote the manuscript. All authors read, corrected, and approved the final manuscript.

## Competing interests

Authors declare no competing interests.

## Supplementary Information

**Figure S1.**
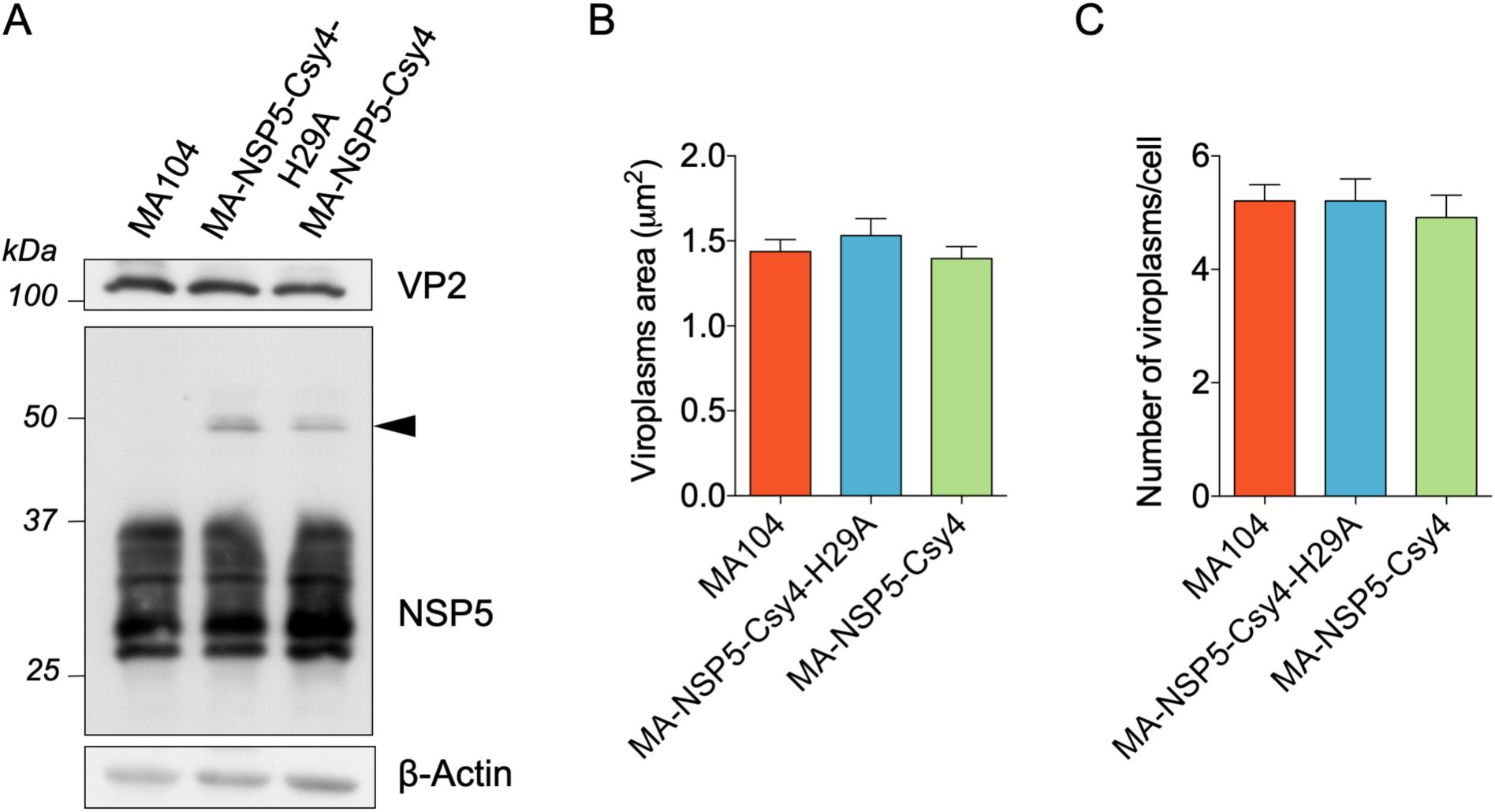
Expression of NSP5-Csy4 fusion proteins did not affect viroplasms formation and viral protein production. A) Western blot of viral proteins VP2 and NSP5 in total cell extracts from with wild-type MA104, MA-NSP5-Csy4, and MA-NSP5-Csy4-H29A cells infected with rRV at a MOI of 5. Black arrowhead indicates NSP5-Csy4 fusions. β-actin was used as loading control. B) Quantification of the size and C) mean number of viroplasms per cell, in the indicated cells lines infected with rRV-wt. Data are expressed as means +/- SEM (n=3).

**Figure S2.**
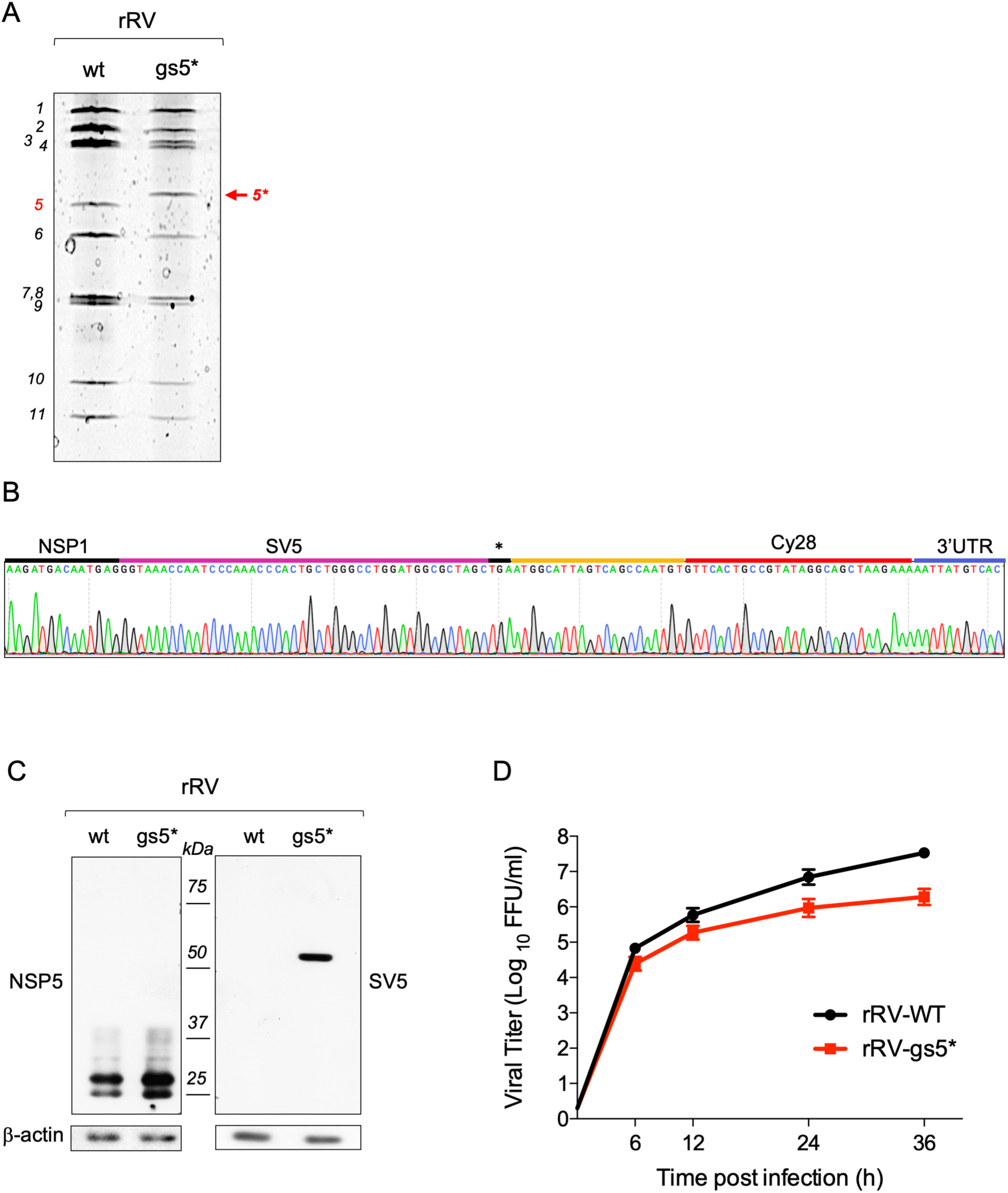
Engineered rRV-gs5*. A) Electrophoretic pattern of dsRNA genome segments of rRV-gs5* and rRV-wt strains grown in MA104 cells at 16 hpi. The number of each genomic segment is indicated on the left. The engineered gs5* of 1704 nt is indicated by the red arrow. B) Chromatogram showing the sequence comprising the modified region of gs5*. C) Western blot of NSP5 and NSP1-SV5 in cell extracts of MA104 cells infected for 5 hours with the rRV-wt or rRV-gs5* strains. β-actin was used as loading control. D) Replication kinetics of rRV-wt and rRV-gs5* infecting MA104 cells. Data are expressed as means +/- SEM (n=3).

**Figure S3.**
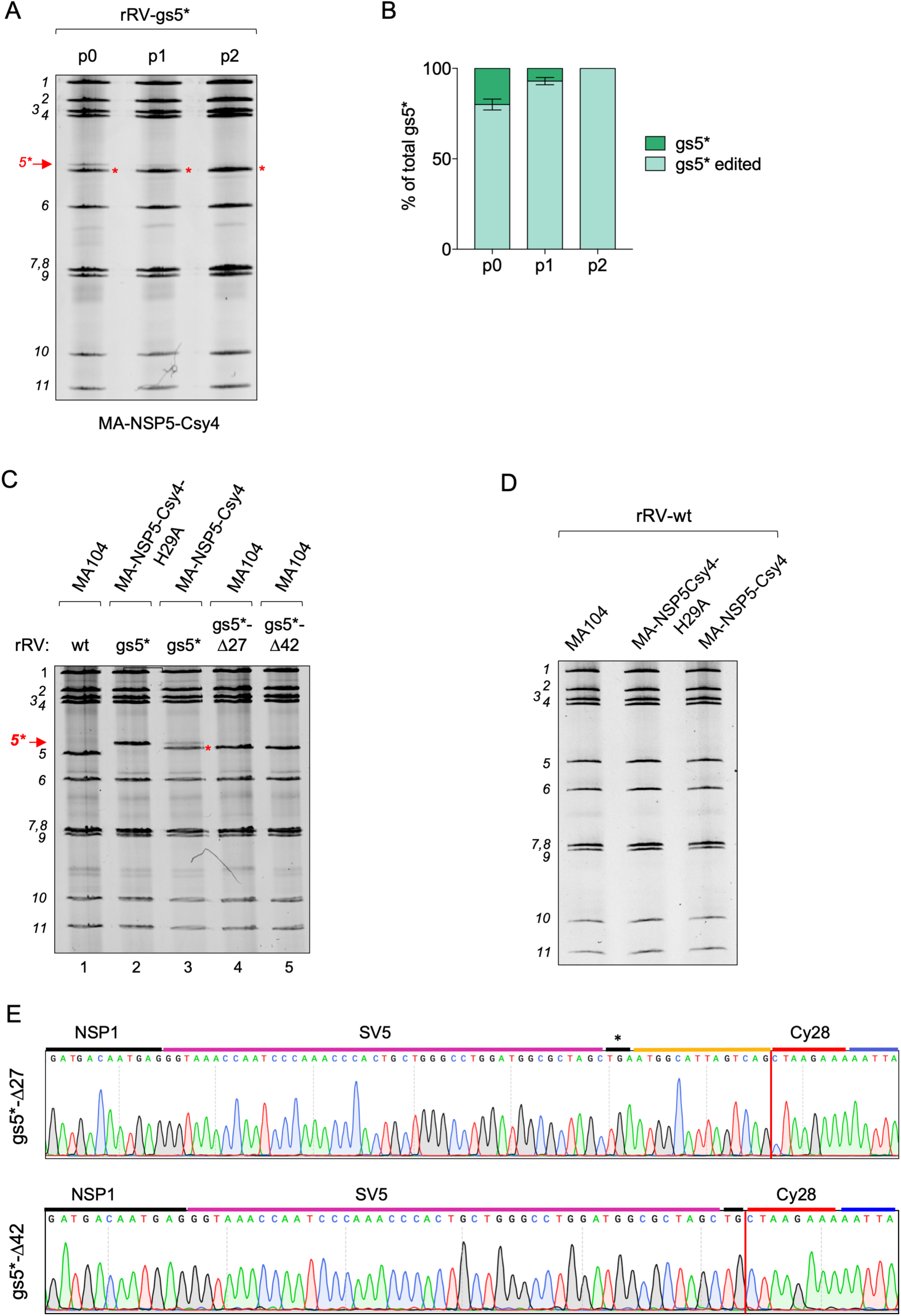
Csy4-mediated editing of rRV-gs5* after multiple infection cycles and isolation of the edited viral strains. A-B) dsRNA electrophoretic migration pattern (A) and quantification of gs5* editing (B) of virus progeny recovered from MA-NSP5-Csy4 cells infected with rRV-gs5* (p0) and after successive passages (p1, p2) in the MA-NSP5-Csy4 cell line. Data are expressed as means +/- SEM (n=3). Red asterisks indicate edited gs5* C) Representative electrophoretic dsRNA migration pattern of the non-edited (lane 2) and edited (lane 3) rRV-gs5* obtained from the indicated cell lines and isolates of the rRV-gs5*-Δ27 (lane 4) and gs5*-Δ42 (lane 5) after six consecutive passages in MA104 cells. Arrow indicates the non-edited gs5*, while red asterisk indicates the edited forms of gs5*. rRV-wt (lane 1) is provided as reference. In A), C) the number of each genomic segment is indicated on the left. D) dsRNA electrophoretic migration of genome segments from rRV-wt replicated in the indicated cells. E) Sequence of gs5*-Δ27 (upper panel) and gs5*-Δ42 (lower panel) after six passages in MA104 cells. Red line indicates the junction.

**Figure S4.**
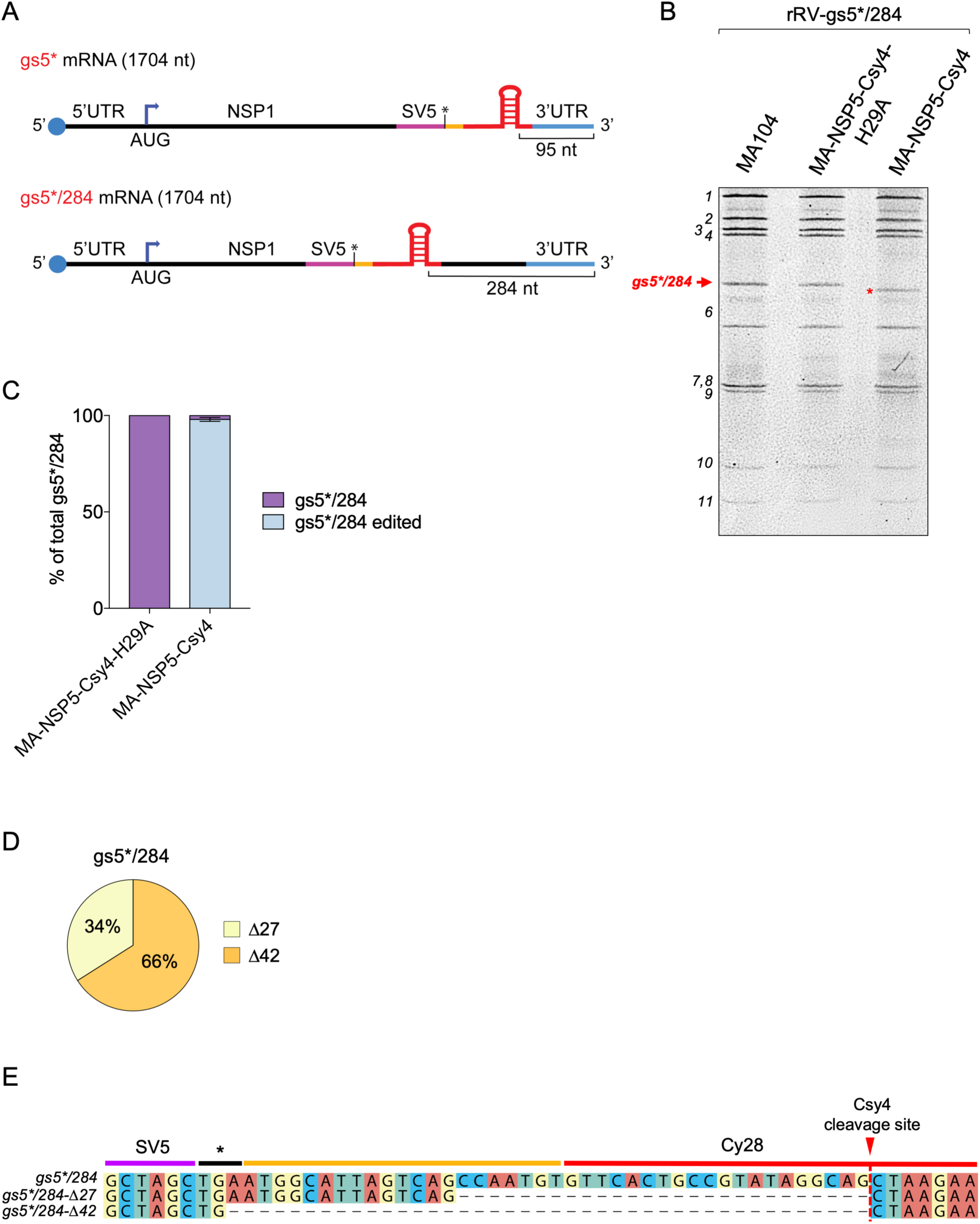
Csy4-mediated editing of rRVgs5*/284. A) Scheme of the modified gs5*/284 and comparison with gs5*. gs5*/284 has the same Cy28 sequence of gs5* inserted 284 nucleotides upstream of the 3’ end (lower panel), instead of 95 nucleotides as in gs5* (upper panel). B) Representative dsRNA electropherotype of rRV-gs5*/284 strain infecting the indicated cell lines at 16 hpi. The number of genomic segments is indicated on the left, while the red asterisk indicates the edited gs5*/284. C) Quantification analysis of the total gs5*/284 shown in the dsRNA electropherotype of (B). Data are expressed as means +/- SEM (n=3). D) Frequency of gs5*/284-Δ27 and gs5*/284-Δ42 sequences obtained from rRV-gs5*/284-infected MA-NSP5-Csy4 cells (22 clones). E) Sequences of the deletion events described in D).

**Figure S5.**
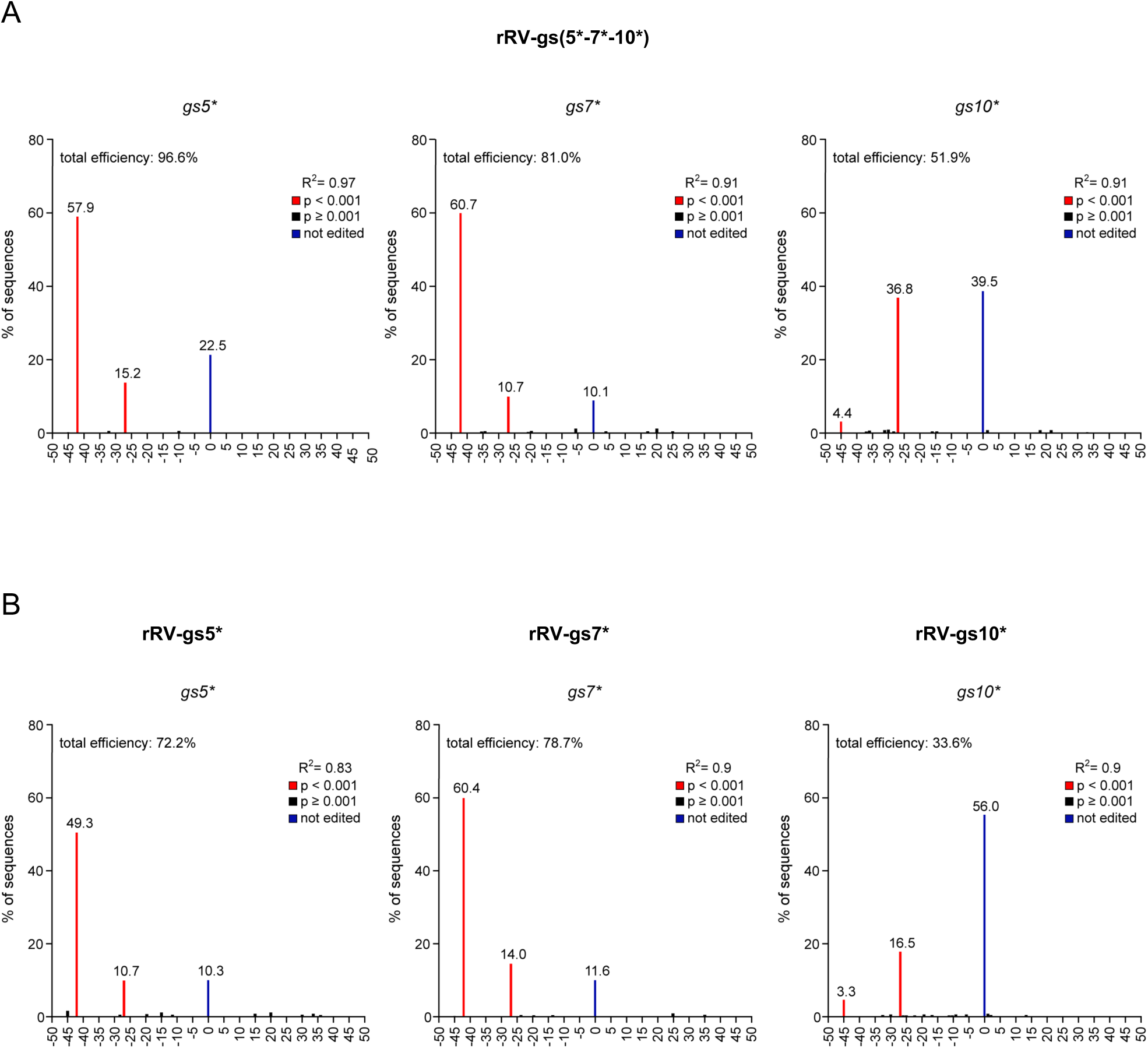
Editing events of multiplexed or individual Csy4-mediated RV genome editing. Indels spectrum by TIDE analysis of the indicated RV genome segments from MA-NSP5-Csy4 cells infected with rRV-gs(5*-7*-10*) (A), or rRV-gs5*, rRV-gs7* and rRV-gs10* (B), representative of n=2 independent experiments.

**Figure S6.**
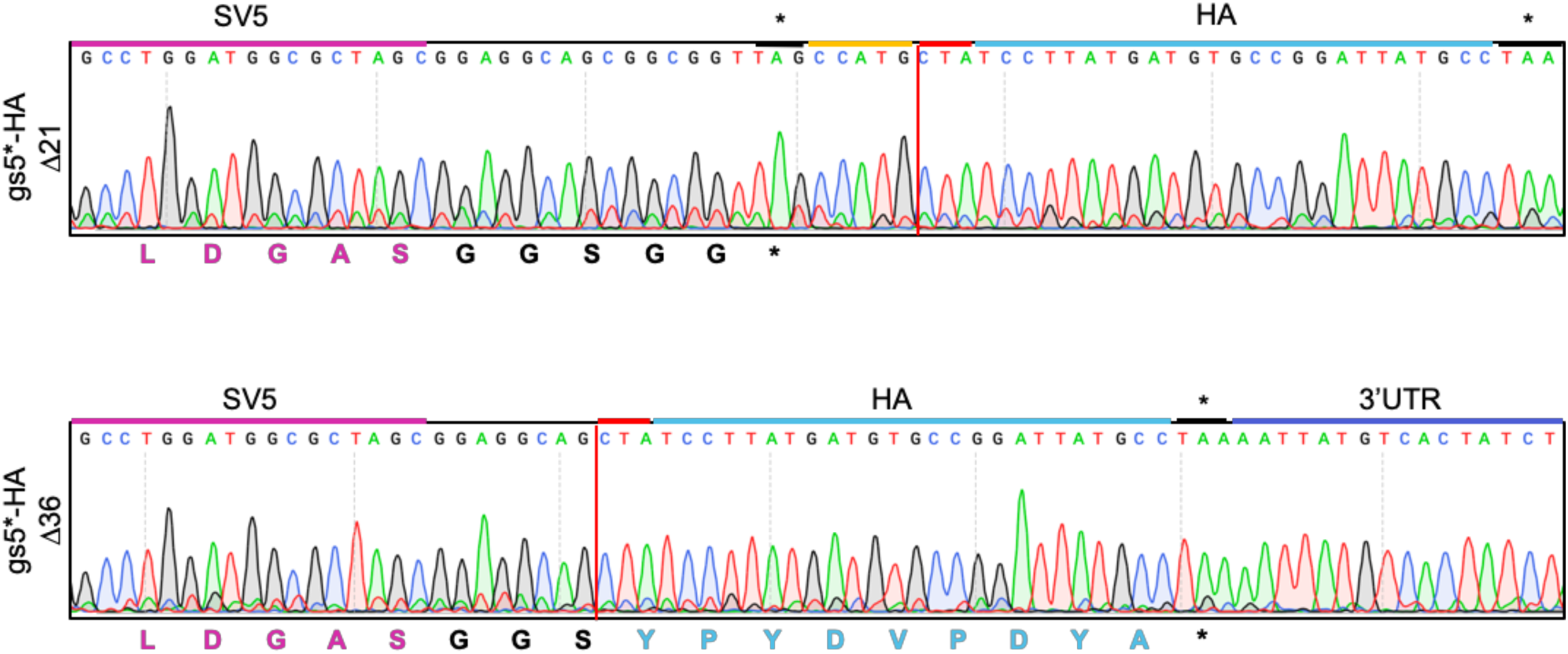
Sequence of gs5*-HA edited segments. Sequence of gs5*-HA-Δ21 (upper panel) and gs5*-HA-Δ36 (lower panel) after a single round of infection in MA-NSP5-Csy4 cells. Amino acid sequence is showed below.

**Figure S7.**
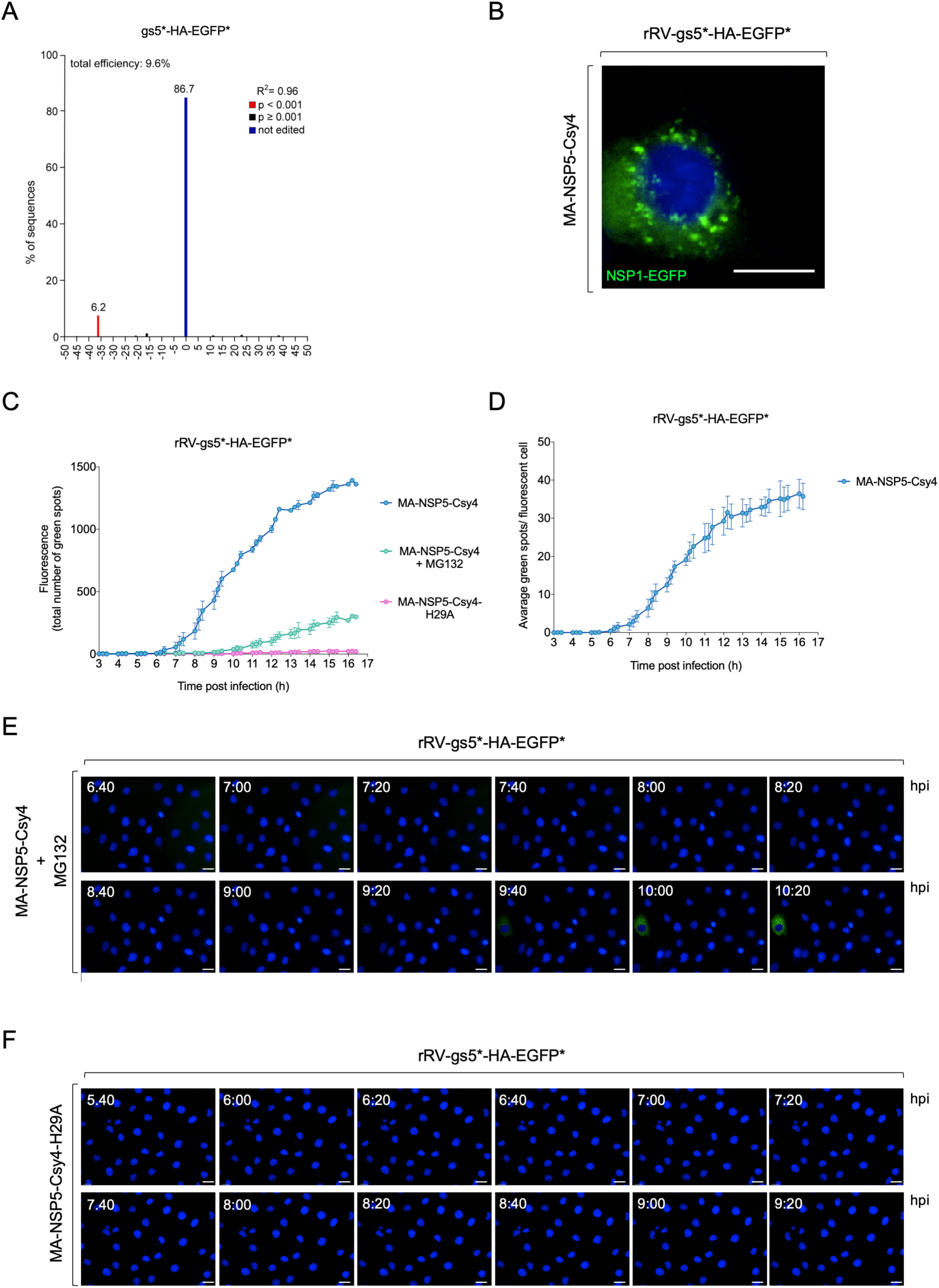
Live-cell imaging of RV secondary transcription and translation. A) Tracking of indels by decomposition (TIDE) of the gs5*-HA-EGFP* genome segment from MA-NSP5-Csy4 cells infected with rRV-gs5*-HA-EGFP*. B) Micrograph of rRV-gs5*-HA-EGFP* infecting MA-NSP5-Csy4. As reported for wild type NSP1 (Murphy and Arnold, 2019), NSP1-EGFP (green) formed spots in infected cells. DAPI is shown in blue. Scale bar 13 μm. C) Total number of NSP1-EGFP dots recorded by live-cell imaging in the indicated gs5*-HA-EGFP* infected cells. D) Average number of NSP1-EGFP dots in NSP1-EGFP positive cell. E-F) Time course of EGFP fluorescence single field micrographs of rRV-gs5*-HA-EGFP* infected MA-NSP5-Csy4 cells treated with MG132 (E) or MA-NSP5-Csy4-H29A (F). Scale bar 50 μm.

**Supplementary Movie 1. Live-cell imaging of EGFP produced by edited RV genomes.** EGFP expression in rRV-gs5*-HA-EGFP* infecting MA-NSP5-Csy4 from experiments as in Figure 7F.

